# Low-density lipoprotein receptor-related protein 1 (LRP1) is a negative regulator of oligodendrocyte progenitor cell differentiation in the adult mouse brain

**DOI:** 10.1101/2020.05.21.108209

**Authors:** Loic Auderset, Kimberley A Pitman, Carlie L Cullen, Renee E Pepper, Bruce V Taylor, Lisa Foa, Kaylene M Young

## Abstract

Low-density lipoprotein receptor-related protein 1 (LRP1) is a large, endocytic cell surface receptor that is highly expressed by oligodendrocyte progenitor cells (OPCs), and LRP1 expression is rapidly downregulated as OPCs differentiate into oligodendrocytes (OLs). We report that the conditional deletion of *Lrp1* from adult mouse OPCs (*Pdgfrα-CreER :: Lrp1^fl/fl^*) increases the number of new myelinating OLs added to brain, but that each new cell elaborates a normal quantity of myelin. OPC proliferation is also elevated following *Lrp1* deletion *in vivo*, however, this is likely to be a secondary, homeostatic response to increased OPC differentiation, as our *in vitro* experiments show that LRP1 is a direct negative regulator of OPC differentiation, not proliferation. Deleting *Lrp1* from adult OPCs also enhances remyelination, as cuprizone-induced lesions are smaller in *Lrp1*-deleted mice, and parenchymal OPCs produce a larger number of mature OLs. These data suggest that the selective blockade of LRP1 function on adult OPCs may enhance myelin repair in demyelinating diseases, such as multiple sclerosis.

## Introduction

Oligodendrocytes (OLs) myelinate the central nervous system (CNS) to facilitate the saltatory conduction of action potentials and provide essential metabolic support to axons (reviewed by 1). The majority of OLs are produced during development, however, new OLs are continuously produced throughout life from oligodendrocyte progenitor cells (OPCs; 2-8), and add new myelin internodes to the CNS (7,9). A number of signaling pathways have been identified that regulate developmental and adult OPC behavior and oligodendrogenesis, including Notch1 (10–12), fibroblast growth factor 2 (13–15), mammalian target of rapamycin (16–18) and platelet-derived growth factor A (19–21) signaling. However, microarray (22) and RNA sequencing (23,24) experiments have uncovered a number of genes that are differentially expressed across OL development, but have no known regulatory function in this lineage. One such gene is the *low-density lipoprotein receptor related protein 1* (*Lrp1*).

LRP1, also known as CD91 or the α2 macroglobulin (α2M) receptor, is highly expressed by OPCs and is rapidly downregulated during OL differentiation (25). This large cell surface receptor, comprising a 515kDa extracellular α-chain and an 85kDa β-chain, could influence OPC behavior in a number of ways, as it interacts with a large variety of ligands, as well as extracellular and intracellular proteins, to facilitate signal transduction (reviewed by 26,27). In other cell types, LRP1 acts as a receptor or coreceptor to initiate intracellular signal transduction, but also facilitates ligand endocytosis, transcytosis or processing (28–34), as well as receptor, channel and transporter trafficking (28,35–40) to influence blood brain barrier permeability (41), lipid metabolism, glucose homeostasis, neuroinflammation (42,43 and reviewed by 44) and synaptic plasticity (45).

*Lrp1* knockout mice are embryonic lethal, as the blastocysts fail to implant (46), but the conditional deletion of *Lrp1* from cultured mouse neural stem and progenitor cells (NSPCs) has been shown to impair NSPC proliferation and particularly reduce the number of OL lineage cells produced (47,48). Furthermore, the conditional deletion of *Lrp1* from *Olig2*^+^ cells (*Olig2-Cre :: Lrp1^fl/fl^* mice) impairs oligodendrogenesis in the developing mouse optic nerve, reducing both the proportion of axons that are myelinated and myelin thickness by postnatal day (P)21 (49). This phenotype may reflect a developmental delay in myelination, as myelin thickness is normal in the optic nerve of *Olig1-Cre :: Lrp1^fl/fl^* mice at P60 (50).

As OPC physiology changes considerably between development and adulthood and can also differ between CNS regions (51–53), we aimed to determine the importance of LRP1 for adult OPC function. The conditional deletion of *Lrp1* from OPCs (*Pdgfrα-CreER :: Lrp1^fl/fl^*) revealed that LRP1 is a negative regulator of adult oligodendrogenesis in the healthy mouse brain. *Lrp1* deletion was associated with an increase in adult OPC proliferation and a significant increase in the number of newborn myelinating oligodendrocytes added to the cortex and corpus callosum. Furthermore, *Lrp1* deletion prior to cuprizone delivery was associated with smaller callosal lesions and a larger number of mature OLs being produced by parenchymal OPCs.

## Materials and Methods

### Animal housing and mice

All animal experiments were approved by the University of Tasmania Animal Ethics (A0016151) and Institutional Biosafety Committees and were carried out in accordance with the Australian code of practice for the care and use of animals for scientific purposes. *Pdgfrα-CreER^T2^* mice (2) were a kind gift from Prof William D Richardson (University College London). *Pdgfrα-CreER^TM^* (5; RRID:IMSR_JAX:018280), *Pdgfrα-H2BGFP* [*Pdgfrα-histGFP* (54);1 RRID:IMSR_JAX:007669] and *Lrp1^fl/fl^* (46; RRID:IMSR_JAX:012604) mice were purchased from Jackson Laboratories. Cre-sensitive *Rosa26-YFP* (55; RRID: IMSR_JAX:006148) and *Tau-mGFP* (56; RRID:IMSR_JAX021162) reporter mice were also purchased from Jackson laboratories. Mice were maintained on a C57BL/6 background and inter-crossed to generate male and female offspring for experimental use. All mice were weaned >P30 to ensure appropriate myelin development, were group housed with same-sex littermates in Optimice micro-isolator cages (Animal Care Systems, Colorado, USA), and were maintained on a 12-hour light / dark cycle at 20°C, with uninhibited access to food and water.

Please note that two distinct *Pdgfrα-CreER* transgenic mouse lines were used in this study: the *Pdgfrα-CreER^TM^* transgenic mouse line (5), was used for the majority of experiments, and the lower efficiency (LE) *Pdgfrα-CreER^T^*^2^ transgenic mouse line (2), was used to perform the *Tau-mGFP* lineage tracing experiments, as we have previously demonstrated that the *Pdgfrα-CreER^TM^* transgenic mouse line cannot be used to induce OPC-specific recombination of the *Tau-mGFP* reporter, despite achieving the OPC-specific recombination of other transgenes (52).

### Genomic DNA extraction and PCR amplification

For genotyping, ear biopsies were digested overnight in DNA extraction buffer (100 mM Tris-HCl, 5 mM EDTA, 200 mM NaCl, 0.2% SDS and 120 ng of proteinase k) at 55°C. Cellular and histone proteins were precipitated by incubating samples with 6M ammonium acetate (Sigma; A1542) on ice, and the DNA subsequently precipitated from the supernatant by incubating with isopropyl alcohol (Sigma; I9516). The DNA pellet was washed in 70% ethanol (Sigma; E7023), resuspended in sterile MilliQ water and used as template DNA for polymerase chain reaction (PCR). Each 25 μL reaction contained: 50-100 ng DNA; 0.5 μL of each primer (100 nmol/mL, GeneWorks); 12.5 μL of GoTaq green master mix (Promega) and MilliQ water. The following primers were used: *Lrp1 5’* CATAC CCTCT CAAACC CCTT CCTG and *Lrp1 3’* GCAAG CTCC CTGCTCA GACC TGGA; *Rosa26 wildtype 5’* AAAGT CGCTC TGAGT TGTTAT, *Rosa26 wildtype 3’* GGAGC GGGAG AAATG GATATG and *Rosa26 mutant 5’* GCGAA GAGTT TGTCC TCAACC; *Cre 5’* CAGGT CTCAG GAGCT ATGTC CAATT TACTG ACCGTA and *Cre 3’* GGTGT TATAAG CAATCC CCAGAA, or *GFP 5’* CCCTG AAGTTC ATCTG CACCAC and *GFP 3’* TTCTC GTTGG GGTCT TTGCTC in a program of: 94°C for 4 min, and 34 cycles of 94°C for 30”, 60°C for 45” (37 cycles for *Rosa26-YFP* genotyping), and 72°C for 60”, followed by 72°C for 10 min. Following gel electrophoresis [1% (w/v) agarose in TAE containing SYBR-safe (ThermoFisher)] the DNA products were visualized using an Image Station 4000M PRO gel system running Carestream software.

### Tamoxifen preparation and administration

Tamoxifen (Sigma) was dissolved in corn oil (Sigma) at a concentration of 40 mg/ml by sonication for 2 hours at 21 °C. Adult mice received tamoxifen (300 mg/kg) daily by oral gavage for 4 consecutive days.

### Tissue preparation and immunohistochemistry

Mice were terminally anaesthetized with an intraperitoneal (i.p) injection of sodium pentobarbital (30mg/kg, Ilium) and were transcardially perfused with 4% (w/v) paraformaldehyde (PFA; Sigma) in phosphate buffered saline (PBS). Brains were cut into 2 mm-thick coronal slices using a 1 mm brain matrix (Kent Scientific) before being post-fixed in 4% (w/v) PFA in PBS at 21°C for 90 min. Tissue was cryoprotected in 20% sucrose (Sigma) in PBS and transferred to OCT (ThermoFisher) before being snap frozen in liquid nitrogen and stored at −80°C.

30 μm coronal brain cryosections were collected and processed as floating sections (as per 57). Cryosections were exposed to primary antibodies diluted in blocking solution [10% (v/v) fetal calf serum (FCS, Serana) and 0.05% (v/v) triton x100 in PBS] and incubated overnight at 4°C on an orbital shaker. Primary antibodies included: rabbit anti-LRP1 (1:500, Abcam ab92544; RRID:AB_2234877); goat anti-PDGFRα (1:100, R&D Systems AF1062; RRID:AB_2236879); rabbit anti-ASPA (1:200, Abcam ab97454; RRID:AB_10679051); rabbit anti-LRP2 (1:100, Abcam ab76969, RRID:AB_10673466); rat anti-GFP (1:2000, Nacalai tesque 04404-26; RRID:AB_2314545); rat anti-MBP (1:100, Millipore MAB386; RRID:AB_94975), rabbit anti-OLIG2 (1:400, Abcam ab9610; RRID:AB_570666); guinea pig anti-IBA1 (1:250, Synaptic Systems 234004; RRID:AB_2493179), and mouse anti-NaBC1 (BCAS1; 1:200, Santa Cruz sc-136342; RRID:AB_10839529).

### EdU administration and labelling

For the *in vivo* labelling of dividing cells, 5-Ethynly-2’-deoxyuridine (EdU; E10415, Thermofisher) was administered to mice via their drinking water at a concentration of 0.2 mg/ml for up to 21 consecutive days (as per 58). For *in vitro* labelling, cells were exposed to 2.5 μg/ml EdU in complete OPC medium (see below) for 10 hours before the cells were fixed with 4% (w/v) PFA in PBS for 15 min at 21°C. The EdU developing cocktail was prepared according to the AlexaFluor-647 Click-IT EdU kit (Invitrogen) instructions, and brain slices were exposed to the developing reagent for 45 min at 21°C, while coverslips of cultured cells were exposed for 15 min. EdU developing was performed immediately after the secondary antibody was washed from tissue or cells during immunohistochemistry or immunocytochemistry.

### Primary OPC culture and *in vitro* gene deletion

The cortices of P1-10 mice were dissected into Earle’s Buffered Salt Solution (EBSS; Invitrogen, 14155-063), diced into pieces ~1 mm^3^ and digested in 0.06 mg/ml trypsin (Sigma, T4799) in EBSS at 37°C for 10 min. The trypsin was inactivated by the addition of FCS, before the tissue was resuspended and triturated in EBSS containing 0.12 mg/ml DNAseI (Sigma, 5025). The cell preparation was filtered through a 40μm sieve (Corning, 352340), centrifuged and resuspended in complete OPC medium [20 ng/ml human PDGF-AA (Peprotech), 10 ng/ml basic fibroblast growth factor (R&D Systems), 10 ng/ml human ciliary neurotrophic factor (Peprotech), 5 μg/ml N-acetyl cysteine (Sigma), 1ng/ml neurotrophin-3 (Peprotech), 1 ng/ml biotin (Sigma), 10μM forskolin (Sigma), 1x penicillin / streptomycin (Invitrogen), 2% B27 (Invitrogen), 50 μg/ml insulin (Sigma), 600 ng/ml progesterone (Sigma), 1 mg/ml transferrin (Sigma), 1 mg/ml BSA (Sigma), 400 ng/ml sodium selenite (Sigma) and 160 μg/ml putrescine (Sigma) in DMEM+ Glutamax (Invitrogen)]. Cells were plated into 6 well plates coated with >300,000 MW Poly D Lysine (PDL; Sigma, P7405). After 7 DIV, the cells were dislodged by incubating in 1:5 TrypLE (Gibco) in EBSS for ~10 min at 37°C, before the trypsin was inactivated by the addition of FBS, and cells were collected into EBSS. OPCs were then purified by immunopanning as previously described (59). In brief, the cell suspension was transferred to a petri dish pre-coated with anti-PDGFRα (BD Pharmigen 558774; RRID:AB_397117) and the OPCs allowed to adhere for 45 min at 21°C. The non-adherent cells were then removed by rinsing with EBSS and the purified OPCs were stripped by treating with TypeLE diluted 1:5 with EBSS for 5 minutes in an incubator. The recovered cells were then plated onto 13mm glass coverslips in complete OPC medium.

For experiments where *Lrp1* was deleted *in vitro*, OPCs were plated in complete OPC medium at a density of 20,000 cells per PDL-treated 13 mm coverslip and allowed to settle for 2 days. OPCs were then exposed to 1μM TAT-Cre (Excellgen, EG-1001) in complete OPC medium at 37°C / 5% CO_2_ for 90 min. The TAT-Cre-containing medium was then removed and replaced with fresh complete OPC medium and the cells returned to the incubator for 48 hours. To induce differentiation, the complete OPC medium was removed and replaced with OPC differentiation medium [complete OPC medium lacking PDGF-AA and containing 4μg/ml triiodothyronine (Sigma)] for 4 days before cells were fixed by exposure to 4% PFA (w/v) in PBS for 15 min at 21°C.

### Whole cell patch clamp electrophysiology

Acute coronal brain slices (300μm) were generated from adult mice carrying the *Pdgfrα-histGFP* transgene, using a VT1200s vibratome (Leica) as previously described (52). Brain slices were transferred to a bath constantly perfused (2 ml/min) with ~21°C artificial cerebral spinal fluid (ASCF) containing: 119 mM NaCl, 1.6 mM KCl, 1 mM NaH_2_PO_4_, 26.2 mM NaHCO_3_, 1.4 mM MgCl_2_, 2.4 mM CaCl_2_, and 11 mM glucose (300 ± 5 mOsm/kg), saturated with 95% O_2_ / 5% CO_2_, Whole cell patch clamp recordings of GFP^+^ cells in the motor cortex were collected using a HEKA Patch Clamp EPC800 amplifier and pCLAMP 10.5 software (Molecular devices; RRID: SCR_011323).

To record AMPA (α-amino-3-hydroxy-5-methyl-4-isoxazolepropionic acid) / kainate receptor currents, recording electrodes (3–6 MΩ) were filled with an internal solution containing: 125 mM Cs-methanesulfonate, 5mM TEA-Cl, 2mM MgCl_2_, 8mM HEPES, 9mM EGTA, 10 mM phosphocreatine, 5 mM MgATP, and 1 mM Na2GTP, and set to a pH of 7.2 with CsOH and an osmolarity of 290 ± 5 mOsm/kg. Upon breakthrough, cells were held at −50 mV and a series of voltage steps (up to +30 mV) applied to determine the presence of a voltage-gated sodium channel current. GFP^+^ cells with a voltage-gated sodium current > 100 pA were considered OPCs. All subsequent recordings were undertaken in ACSF containing 50 μm (2R)-amino-5-phosphonovaleric acid (APV; Sigma) and 1 μM tetrodotoxin (TTX; Sigma). Cells were held at −60 mV and currents elicited by applying 200 ms voltage steps from −80 to 20 mV (20 mV increments). After taking baseline recordings, currents were then elicited in ACSF containing 100 μM kainate. The mean steady state current (last 100 ms) of each voltage step was measured.

Voltage gated calcium channel (VGCC) current recordings were made using solutions previously described (52). All other voltage gated currents (potassium and sodium) were blocked. To record L-type VGCC currents, OPCs were held at −50 mV and a series of 500 ms voltage steps (−60 to +30 mV) applied using a P/N subtraction protocol. The current-density relationship is presented as the average steady state current (the last 100ms of the voltage steps) from ~3 recordings per cell. To elicit currents through T-type VGCCs, OPCs were held at −50 mV and the cell was hyperpolarized to −120 mV for 200 ms before applying voltage steps from −70 mV to 30 mV (as per 60,61). The maximum amplitude of the fast, transient inward current, revealed by the brief hyperpolarization, was measured from ~3 recordings per cell.

Access resistance was measured before and after all recordings and an access resistance >20 MΩ resulted in exclusion of that recording. Due to the high membrane resistance of OPCs (>1 GΩ) during VGCC current recordings, recordings were made without series resistance compensation. However, series resistance compensation was applied for AMPA current recordings (60-80%). Measurements were made from each data file using Clampfit 10.5.

### Cuprizone administration and black-gold myelin staining

Mice were transferred onto a diet of crushed mouse food (Barrestock) containing 0.2% (w/w) cuprizone powder (C9012, Sigma), which was refreshed every 2 days for 5 weeks. Mice were perfusion fixed and their tissue processed as described previously, and 30μm coronal brain floating cryosections collected into PBS. Cryosections were transferred onto glass microscope slides (Superfrost) and allowed to dry, before being rehydrated in milliQ water for 3 min and incubated with preheated 0.3% black-gold II stain (Millipore, AG105) at 60°C for 30 min. Slides were washed twice with milliQ water before being incubated with preheated 1% (v/v) sodium thiosulphate solution at 60°C for 3 min, washed in milliQ water (3×2 min), dehydrated using a series of graded alcohol steps, incubated in xylene (Sigma, 214736) for 3 min, and mounted with DPX mounting medium (Sigma, 06522).

### Microscopy and statistical analyses

Fluorescent labelling was visualized using an UltraView Spinning Disc confocal microscope with Volocity software (Perkin Elmer, Waltham, USA). The motor cortex and corpus callosum were defined as regions of interest using anatomical markers identified in the Allen Mouse Brain Atlas, in brain sections collected between Bregma level 1.10 mm and −0.10 mm. Confocal images were collected using standard excitation and emission filters for DAPI, FITC (Alexa Fluor-488), TRITC (Alexa Fluor-568) and CY5 (Alexa Fluor-647). To quantify cell density, or the proportion of cells that proliferate or differentiate, a 10x or 20x air objective was used to collect images with 2μm z-spacing, that spanned the defined region of interest within a brain section, and these images were stitched together using Volocity software to create a single image of that region for analysis. A minimum of 3 brain sections were imaged per mouse. To quantify oligodendrocyte morphology and measure myelin internodes, a 40x (air) or 60x (water) objective was used to collect images with 1 μm z-spacing of individual mGFP^+^ OLs or single fields of view containing internodes within each region of interest. Black-gold myelin staining was imaged using a light microscope with a 2.5x objective, and images were manually stitched together using Adobe Photoshop CS6 to recreate the region of interest. Cell counts were performed by manually evaluating the labelling of individual cells, and area measurements were made by manually defining the region of interest within Photoshop CS6 (Adobe, San Jose, USA) or Image J (NIH, Bethesda, Maryland). All measurements were made blind to the experimental group and treatment conditions.

Statistical comparisons were made using GraphPad Prism 8.0 (La Jolla CA, USA; RRID: SCR_002798). Data were first assessed using the D’Agostino-Pearson normality test. Data that were normally distributed were analysed by a parametric test [t-test, one-way analysis of variance (ANOVA) or two-way ANOVA for group comparisons with a Bonferroni post-hoc test], and data that were not normally distributed were analysed using a Mann-Whitney U test or Kolmogorov-Smirnoff test. Lesion size (black-gold staining) data were analyzed using a t-test with a Welch’s correction, to account for the uneven variance between groups. Data sets with n=3 in any group were analysed using parametric tests, as the non-parametric equivalents rely on ranking and are unreliable for small sample sizes (GraphPad Prism 8.0). To determine the rate at which OPCs become labelled with EdU over time, these data were analysed by performing linear regression analyses. Details of the statistical comparisons are provided in each figure legend or in text when the data are not presented graphically. Statistical significance was defined as p<0.05.

## Results

### LRP1 can be successfully deleted from OPCs in the adult mouse brain

In order to determine the role that LRP1 plays in regulating adult myelination, *Lrp1* was conditionally deleted from OPCs in young adult mice. Tamoxifen was administered to P50 control (*Lrp1^fl/fl^*) and *Lrp1*-deleted (*Pdgfrα-CreER^TM^ :: Lrp1^fl/fl^*) mice and brain tissue examined 7 or 30 days later (at P50+7 and P50+30, respectively). Coronal brain cryosections from control (**Fig. 1A**) and *Lrp1*-deleted mice (**Fig. 1B**) were immunolabelled to detect LRP1 (red) and OPCs (PDGFRα, green). Consistent with our previous findings (25), essentially all OPCs in the corpus callosum of control mice expressed LRP1 (**Fig. 1C**; P50+7: 99% ± 0.6%, mean ± SD for n=4 mice; P50+30: 99.7% ± 0.3%, mean ± SD for n=3 mice). However, in the corpus callosum of P50+7 *Lrp1*-deleted mice, only 2% ± 0.8% of PDGFRα^+^ OPCs expressed LRP1 (mean ± SD for n=4 mice), and by P50+30, only 0.5% ± 0.5 of OPCs expressed LRP1 (mean ± SD for n=3 mice; **Fig. 1C**), confirming the successful deletion of *Lrp1* from adult OPCs. Similarly, in the motor cortex of P50+7 control mice, 100% ± 0% of PDGFRα^+^ OPCs expressed LRP1, while only 0.4% ± 0.4% of PDGFRα^+^ OPCs expressed LRP1 in the motor cortex of *Lrp1*-deleted mice (mean ± SD for n=4 mice per genotype). Furthermore, *Lrp1*-deletion was specific, as other brain cell types that express LRP1, such as neurons and astrocytes, retained their expression of LRP1 (e.g. white arrows in **Fig. 1B**). As the Cre-mediated recombination of the *Lrp1^fl/fl^* transgene deletes the extracellular coding region of *Lrp1*, recombination was also confirmed by performing a PCR analysis of genomic DNA from the brains of control (*Lrp1 ^fl/fl^*) and *Lrp1*-deleted (*Pdgfrα-CreER^TM^ :: Lrp1^fl/fl^*) mice after tamoxifen treatment. *Lrp1*-deletion enabled the amplification of a recombination-specific DNA product from *Lrp1*-deleted brain DNA that was not amplified from control brain DNA (**Fig. 1D**). These data confirm that tamoxifen administration to *Pdgfrα-CreER^TM^ :: Lrp1^fl/fl^* transgenic mice efficiently and specifically deletes *Lrp1* from adult OPCs.

**Figure 1:**
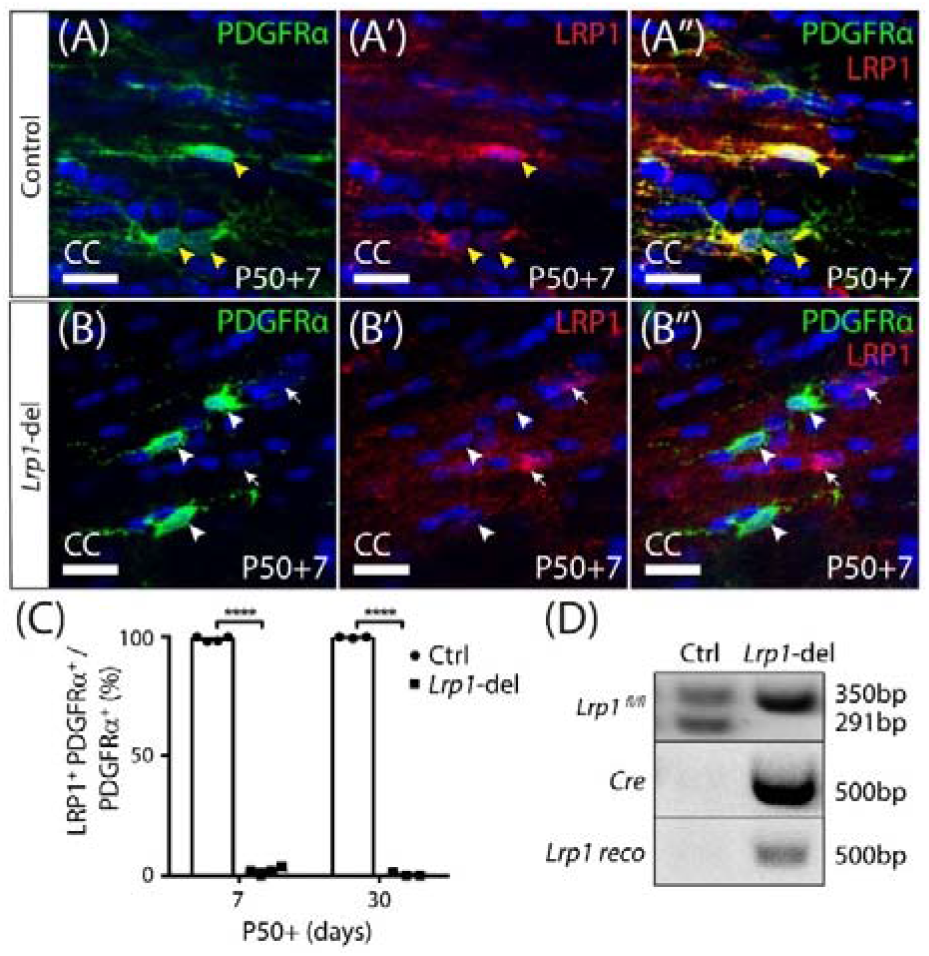
Lrp1 can be conditionally deleted from adult OPCs at high efficiency. Coronal brain sections from P57+7 and P57+30 control (*Pdgfrα-CreER^TM^) and Lrp1*-deleted (*Pdgfrα-CreER^TM^ :: Lrp1^fl/fl^*) mice were immunolabelled to detect OPCs (PDGFRα, green), LRP1 (red) and cell nuclei (Hoescht 33342, blue). (**A-A’’**) Compressed z-stack confocal image of LRP1^+^ OPCs (solid yellow arrow heads) in the corpus callosum (CC) of a P50+7 control mouse. (**B-B’’**) Compressed z-stack confocal image of LRP1-neg OPCs (solid white arrow heads) in the CC of a P50+7 *Lrp1*-deleted mouse. White arrows indicate PDGFR-neg cells that remain LRP1^+^ in the *Lrp1*-deleted mice. (**C**) The proportion (%) of PDGFRα^+^ OPCs that express LRP1 in P50+7 and P50+30 control and *Lrp1*-deleted mice [mean ± SD for n≥3 mice per genotype per time-point; 2-way ANOVA: *Genotype F* (1, 10) = 2.8, p <0.0001; *Time F* (1, 10) = 0.52, p = 0.5; *Interaction F* (1, 10) = 3.44, p = 0.09]. Bonferroni multiple comparisons **** p ≤ 0.0001. (**D**) PCR amplification of genomic DNA from the brain of P50+7 control (*Pdgfrα-CreER^TM^*) and *Lrp1*-deleted (*Pdgfrα-CreER^TM^ :: Lrp1^fl/fl^*) mice indicates that recombination (producing the *Lrp1* reco band) only occurs in *Lrp1*-deleted mice. Scale bars represent 17 μm.

### *Lrp1*-deletion increases adult OPC proliferation

OPCs divide more frequently in white than grey matter regions of the adult mouse CNS (62), and their homeostatic proliferation ensures that a stable pool of progenitors is maintained (6). To determine whether LRP1 regulates the frequency at which OPCs re-enter the cell cycle to divide, or the fraction of OPCs that proliferate, we delivered a thymidine analogue, EdU, to P57+7 control and *Lrp1*-deleted mice via their drinking water for 2, 4, 6 or 20 days. Coronal brain cryosections from control (**Fig. 2A-D**) and *Lrp1*-deleted (**Fig. 2E-H**) mice were processed to detect PDGFRα^+^ OPCs (green) and EdU (red). When quantifying the proportion of OPCs that became EdU labelled over time, we found that 20 days of EdU-delivery resulted in EdU uptake by all OPCs in the corpus callosum of control and *Lrp1*-deleted mice (100% ± 0% and 100% ± 0% respectively), indicating that the proportion of OPCs that can proliferate is not influenced by LRP1 signaling. Furthermore, the rate of EdU incorporation by OPCs was equivalent in the corpus callosum of control and *Lrp1*-deleted mice (**Fig. 2I**), suggesting that LRP1 does not influence the rate at which OPCs enter or transition through the cell cycle to become EdU-labelled. While OPCs in the motor cortex incorporated EdU at a slower rate than those in the corpus callosum (compare the slope of the regression lines in **Fig. 2I** and **Fig. 2J**), OPC proliferation in the motor cortex was also unaffected by *Lrp1*-deletion (**Fig. 2J**).

**Figure 2:**
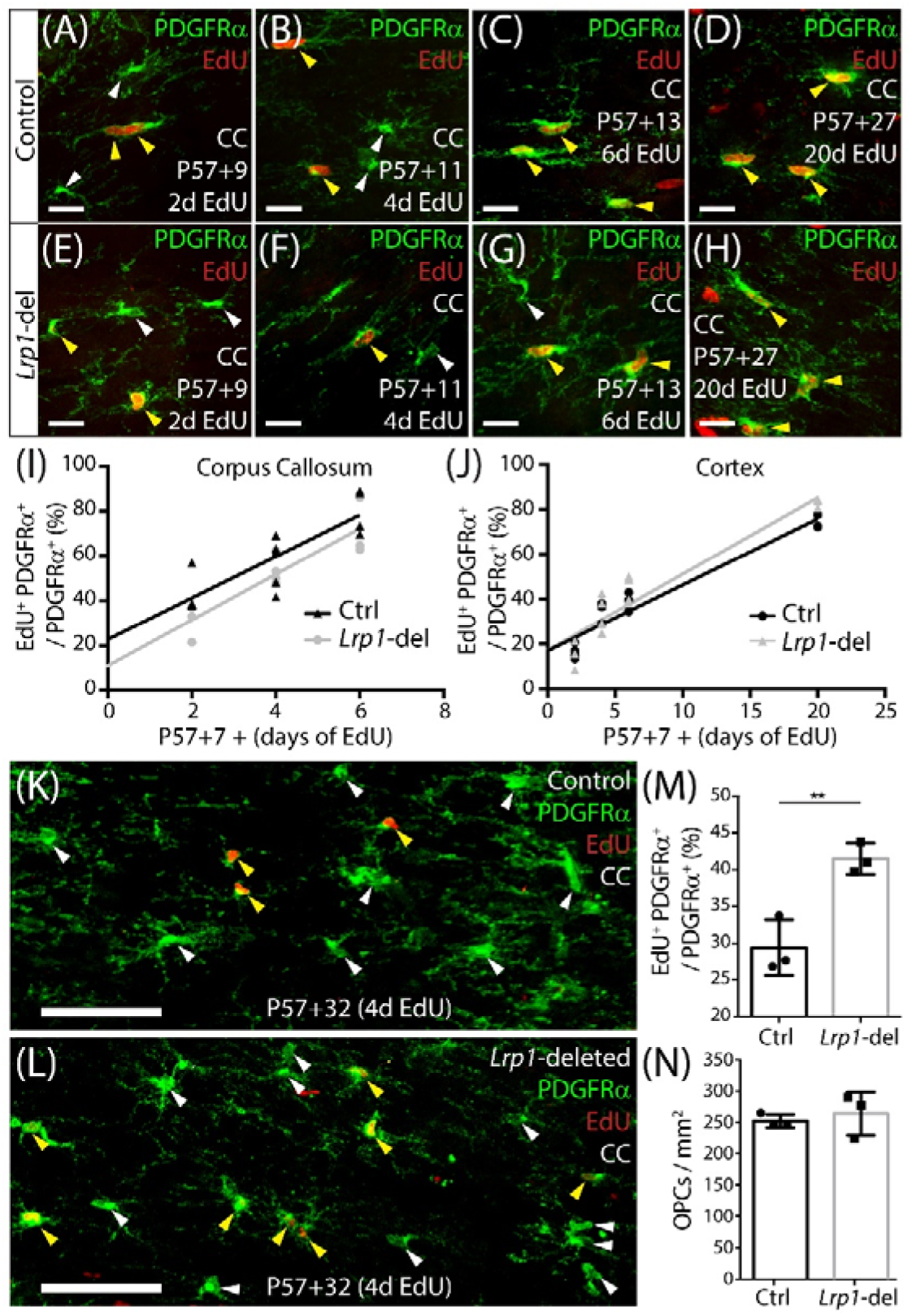
Adult OPC proliferation increases following Lrp1 deletion. (**A-H**) Compressed confocal z-stacks from the corpus callosum (CC) of control (*Pdgfrα-CreER^TM^*) and *Lrp1*-deleted (*Pdgfrα-CreER™:: Lrp1^fl/fl^*) mice, immunolabelled to detect OPCs (PDGFRα, green) and EdU (red) after 2, 4, 6 or 20 days of EdU delivery. (**I**) Graph showing that the proportion (%) of OPCs that incorporate EdU, after 2, 4 or 6 days of delivery, in the CC of control (black) and *Lrp1*-deleted (grey) mice (n ≥ 3 mice per genotype per timepoint). The rate of EdU uptake was unaffected by genotype (p=0.7; linear regression for controls: m = 9.2 ± 1.8 % per day and R^2^ = 0.7; linear regression for *Lrp1*-deleted: m = 10.2 ± 1.8 % per day and R^2^ = 0.8). (**J**) Graph showing that the proportion (%) of OPCs that incorporate EdU, after 2, 4, 6 or 20 days of delivery, in the motor cortex of control (black) and *Lrp1*-deleted (grey) mice (n ≥ 3 mice per genotype per timepoint). The rate of EdU uptake was unaffected by genotype (p=0.3; linear regression for control: m = 2.9 ± 0.3 cells per day and R^2^ = 0.9; linear regression for *Lrp1*-deleted: m = 3.4 ± 0.3 cells per day and R^2^ = 0.9). (**K-L**) Compressed confocal z-stacks from the CC of P57+32 control and *Lrp1*-deleted mice that received EdU via the drinking water for 4 consecutive days (from P57+28), and were immunolabelled to detect OPCs (PDGFRα, green) and EdU (red). (**M**) The proportion (%) of OPCs that were EdU labelled in the CC of P57+32 control mice (black) and *Lrp1*-deleted mice (grey) that received 4 days of EdU labelling (mean ± SD, n=3 mice per genotype; unpaired t-test, p=0.008). (**N**) Quantification of the density of OPCs in the CC of P57+32 control mice (black) and *Lrp1*-deleted mice (grey) (mean ± SD, n=3 mice per genotype; unpaired t-test, p= 0.6). Solid white arrow heads indicate EdU-neg OPCs. Solid yellow arrowheads indicate EdU^+^ OPCs. Scale bars represent 17 μm (A-H) or 70 μm (K, L).

These data indicate that the loss of LRP1 does not immediately influence OPC proliferation, however, LRP1 may regulate processes such as receptor and channel recycling at the cell membrane (36,40,63–65), such that *Lrp1*-deletion may not immediately perturb OPC behavior. To explore this possibility, we delivered tamoxifen to young adult (P57) control and *Lrp1*-deleted mice and waited a further 28 days before administering EdU via the drinking water for 4 consecutive days. Coronal brain cryosections from P57+32 control (**Fig. 2K**) and *Lrp1*-deleted (**Fig. 2L**) mice were processed to detect PDGFRα^+^ OPCs (green) and EdU (red). The proportion of OPCs that incorporated EdU over the 4day labelling period was significantly higher in the corpus callosum of *Lrp1*-deleted mice than controls (**Fig. 2M**). This increase in OPC proliferation was not accompanied by a change in the density of PDGFRαΛ OPCs, which was equivalent in the corpus callosum of control and *Lrp1*-deleted mice (**Fig. 2N**). These data suggest that *Lrp1* deletion from adult OPCs results in a delayed increase in OPC proliferation. As OPC density remains unchanged, the large number of new cells must either differentiate into new OLs or die.

### LRP1 is a negative regulator of adult oligodendrogenesis

To determine whether LRP1 regulates OL production by adult OPCs, tamoxifen was given to P57 control (*Pdgfrα-CreER^TM^ :: Rosa26-YFP*) and *Lrp1*-deleted (*Pdgfrα-CreER^TM^ :: Rosa26-YFP :: Lrp1^fl/fl^*) mice, to fluorescently label adult OPCs and the new OLs they produce. At P57+14, coronal brain cryosections were immunolabeled to detect YFP (green), PDGFRα (red) and OLIG2 (blue), to confirm the specificity of labelling (**Fig. S1**). Consistent with our previous findings in control mice (57), all YFP^+^ cells in the corpus callosum of control and *Lrp1*-deleted mice were either PDGFRα^+^ OLIG2^+^ OPCs or PDGFRα-negative OLIG2^+^ newborn OLs (**Fig. S1**). In the motor cortex, the vast majority of YFP^+^ cells expressed OLIG2 (control: 96.2% ± 0.91; *Lrp1*-deleted: 94.3% ± 1.02, mean ± SD for n=3 mice per genotype; **Fig. S1**), and the small number of YFP^+^ OLIG2-negative cells identified in the cortex had the morphological characteristics of neurons, consistent with previous reports that the *Pdgfrα* promoter is active in a small subset of cortical neurons (58), and were excluded from all subsequent analyses.

To determine whether LRP1 influences oligodendrogenesis, we quantified the proportion of YFP^+^ cells that were PDGFRα-negative OLIG2^+^ newborn OLs in the corpus callosum (**Fig. 3A-F**) or motor cortex (**Fig. 3G-L**) of P57+7, P57+14, P57+30 and P57+45 control and *Lrp1*-deleted mice. At P57+7 and P57+14, oligodendrogenesis was equivalent in the corpus callosum of control and *Lrp1*-deleted mice, however by P57+30, a larger proportion of YFP^+^ cells had become newborn OLs in the corpus callosum of *Lrp1*-deleted mice, and this effect was sustained at P57+45 (**Fig. 3M**). Similarly, for the first two weeks, OL production was equivalent for OPCs in the motor cortex of control and *Lrp1*-deleted mice, however, by P57+30, the proportion of YFP^+^ cells that were newborn OLs was higher in the motor cortex of *Lrp1*-deleted mice than controls (**Fig. 3N**). At P57+30, we also performed cell density measurements and found that that the density of new OLs was significantly increased in the corpus callosum (control: 107.2 ± 14.9 cells / mm^2^; *Lrp1*-del: 161.8 ± 27.4 cells / mm^2^; mean ± SD, n= 7 control and n=4 *Lrp1*-deleted mice; t-test, p=0.0005) and motor cortex (control: 42.55 ± 9.2 cells / mm^2^; *Lrp1*-del: 61.34 ± 7.0 cells / mm^2^; mean ± SD, n=4 mice per genotype; t-test, p=0.004) of *Lrp1*-deleted mice compared to controls. These results suggest that LRP1 is a negative regulator of adult oligodendrogenesis.

**Figure 3:**
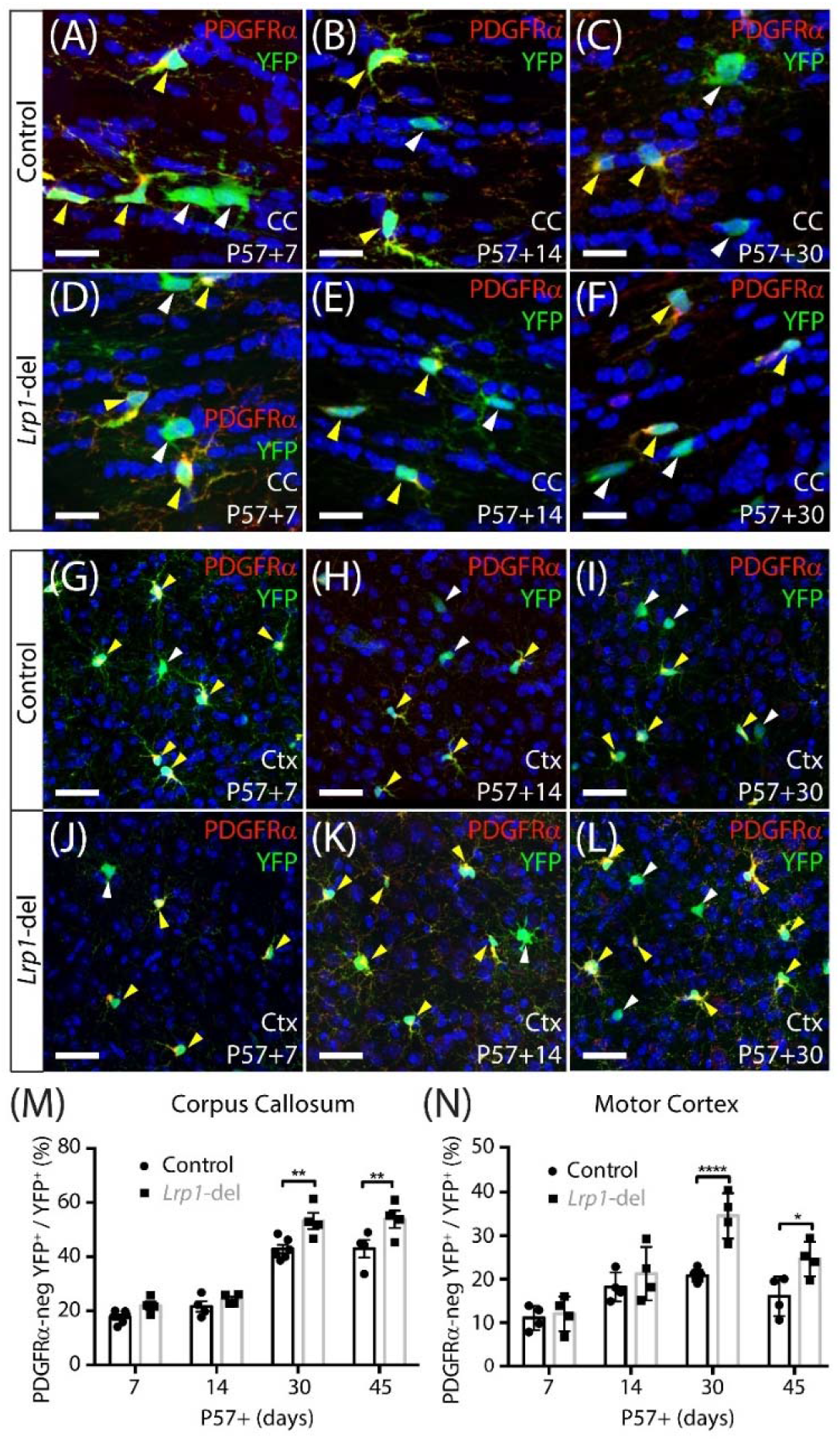
LRP1 reduces oligodendrogenesis in the adult mouse corpus callosum and motor cortex. (**A-L**) Confocal images of the corpus callosum (CC; A-F) and motor cortex (Ctx; G-L) of P57+7, P57+14 and P57+30 control (Pdgfrα-CreER^TM^ :: Rosa26-YFP) and Lrp1-deleted (Pdgfrα-CreER^TM^ :: Rosa26-YFP :: Lrp1^fl/fll^) mice immunolabelled to detect PDGFRα (red), YFP (green) and the nuclear marker Hoescht 33342 (blue). Solid yellow arrowheads indicate YFP^+^ PDGFRα^+^ OPCs. Solid white arrowheads indicate YFP^+^ PDGFRα-neg newborn OLs. (M) Graphical representation of the proportion (%) of YFP^+^ cells that are YFP^+^ PDGFRα-neg OLIG2^+^ newborn OLs in the CC of control and Lrp1-deleted mice [mean ± SD for n≥4 mice per genotype per timepoint; 2-way ANOVA: Genotype F (1, 28) = 22.3, p <0.0001; Time F (3, 28) =109.7, p <0.0001; Interaction F (3, 28)= 1.902, p = 0.15]. (N) Graphical representation of the proportion (%) of YFP^+^ cells that are YFP^+^ PDGFRα-neg OLIG2^+^ newborn OLs in the Ctx of control and Lrp1-deleted mice [mean ± SD for n≥4 mice per genotype per timepoint; 2-way ANOVA: Genotype F (1, 26) = 22.5, p <0.0001; Time F (3, 26) = 23.4, p <0.0001; Interaction F (3, 26) = 4.56, p = 0.011]. Bonferroni multiple comparisons: * p<0.05, ** p<0.01, ****p<0.0001. Scale bars represent 17 μm (A-F) and 34 μm (G-L).

### LRP1 reduces the generation of mature, myelinating oligodendrocytes

As OPCs differentiate, they rapidly downregulate their expression of PDGFRα, the NG2 proteoglycan and voltage-gated sodium channels (NaV) (58,66–69), and become highly ramified pre-myelinating OLs, that either die or continue to mature into myelinating OLs, that are characterized by the elaboration of myelin internodes (4,6,52,62,70,71). In order to determine whether *Lrp1*-deletion increases the number of myelinating OLs, we fluorescently labelled a subset of OPCs in the adult mouse brain with a membrane-targeted form of green fluorescent protein (GFP), allowing us to visualize the full morphology of the OPCs and the OLs they produce. We have previously shown that tamoxifen delivery to adult *Pdgfrα-CreER^TM^* :: *Tau-GFP* mice does not result in the specific fluorescent labelling of OPCs and their progeny (52). Therefore, for this experiment, we instead delivered tamoxifen to adult LE-control (*Pdgfrα-CreER^T2^ :: Tau-GFP*) and LE-*Lrp1*-deleted (*Pdgfrα-CreER^T2^ :: Tau-GFP :: Lrp1^fl/fl^*) mice. The *Pdgfrα-CreER^T2^* transgenic mouse (2) has a lower recombination efficiency (LE) than the *Pdgfrα-CreER^TM^* transgenic mouse (5), so we first evaluated the efficiency of *Lrp1* deletion using this mouse model. Coronal brain cryosections from P57+30 LE-control and LE-*Lrp1*-deleted mice were immunolabelled to detect PDGFRα and LRP1 (**Fig. 4 A, B**), and while 100% ± 0% of PDGFRα^+^ OPCs expressed LRP1 in the motor cortex of LE-control mice, only 35% ± 9% of PDGFRα^+^ OPCs expressed LRP1 in the motor cortex of LE-*Lrp1*-deleted mice (mean ± SD for n=3 mice per genotype). The recombination efficiency was similar in the corpus callosum, with 100% ± 0% of OPCs expressing LRP1 in LE-control mice and only 37% ± 7 in LE-*Lrp1*-deleted mice (**Fig. 4C**).

**Figure 4:**
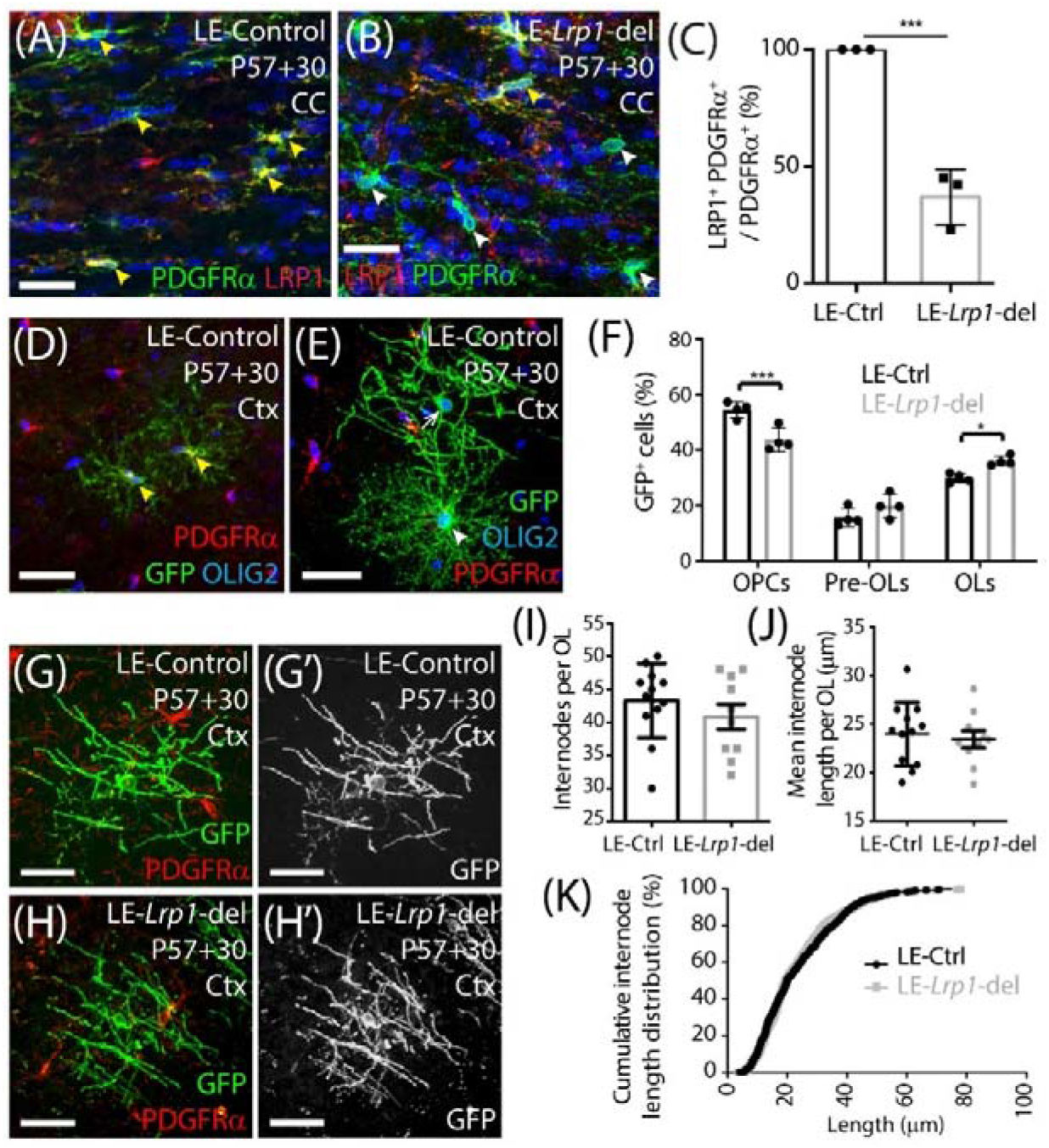
Lrp1-deletion increases the number of mature, myelinating OLs added to the motor cortex of adult mice. (**A-B**) Compressed confocal z-stack images of the corpus callosum (CC) in P57+30 LE-control (*Pdgfrα-CreER^T2^*) and LE-*Lrp1*-deleted (*Pdgfrα-CreER^T2^ :: Lrp1^fl/lf^*) mice immunolabelled to detect OPCs (PDGFRα, green), LRP1 (red) and Hoescht 33342 (blue). Solid yellow arrowheads indicate OPCs that express LRP1. Solid white arrowheads indicate OPCs that do not express LRP1. (**C**) The proportion (%) of PDGFRα^+^ OPCs in the CC of LE-control and the LE-*Lrp1*-del mice that express LRP1 (mean ± SD, n=3 mice per group; unpaired t-test, *** p =0.0008). (**D-E**) Compressed confocal z-stack images from the motor cortex (Ctx) of a P57+30 control (LE-*Pdgfrα-CreER^T2^ :: Tau-mGFP*) mouse immunolabelled to detect PDGFRα (red), GFP (green) and OLIG2 (blue). Solid yellow arrowheads indicate GFP^+^ PDGFRα^+^ OLIG2^+^ OPCs. Solid white arrowhead indicates a GFP^+^ PDGFRα-neg OLIG2^+^ newborn pre-myelinating OL. The white arrow indicates a GFP^+^ PDGFRα-neg OLIG2^+^ newborn myelinating OL. (**F**) Quantification of the proportion (%) of GFP^+^ cells that are PDGFRα^+^ OLIG2^+^ OPCs, PDGFRα-neg OLIG2^+^ premyelinating OLs (pre-OLs) and PDGFRα-neg OLIG2^+^ myelinating OLs (OLs) [mean ± SD for n = 4 mice per genotype; 2-way ANOVA: *Maturation stage F (2, 18) = 195.1, p <0.0001; Genotype F (1, 18) = 0.032, p = 0.85; Interaction F (2, 18)= 17.1, p <0.0001]*. Bonferroni multiple comparisons: * p = 0.046 and *** p = 0.0004. (**G-H**) Compressed z-stack confocal images of GFP^+^ PDGFRα-neg myelinating OLs in the motor cortex of P57+30 LE-control and LE-*Lrp1*-deleted mice. (**I**) The number of internodes elaborated by individual GFP^+^ myelinating OLs in the motor cortex of LE-control and LE-*Lrp1*-deleted mice (mean ± SEM for n ≥ 10 OLs from n=3 mice per genotype; Mann Whitney Test, p = 0.38). (**J**) The average length of internodes elaborated by individual GFP^+^ myelinating OLs in LE-control and LE-*Lrp1*-deleted mice (mean ± SEM for n ≥ 10 OLs from n=3 mice per genotype; unpaired t-test, p= 0.67). (**K**) Cumulative length distribution plot for GFP^+^ internodes measured in the motor cortex of P57+30 LE-control and LE-*Lrp1*-deleted mice (n=519 LE-control GFP^+^ internodes and n=408 LE-*Lrp1*-deleted GFP^+^ internodes measured from n=3 mice per genotype; K-S test, D = 0.053, p= 0.5). Scale bars represent 34 μm (A, B) or 17 μm (G, H).

While only ~65% of OPCs lacked LRP1 in the LE-*Lrp1*-deleted mice, this was sufficient to increase adult oligodendrogenesis. Brain cryosections from P57+30 LE-control and LE-*Lrp1*-deleted mice were immunolabelling to detect GFP (green), PDGFRα (red) and OLIG2 (blue) (**Fig. 4D, E**), and we found that the proportion of GFP^+^ cells that became PDGFRα-negative OLIG2^+^ newborn OLs was significantly elevated in the motor cortex of LE-*Lrp1*-deleted mice (56.3% ± 2.06%) compared to control mice (49.2% ± 1.51, mean ± SD for n=4 mice per genotype; t-test p=0.03). Furthermore, by using the morphology to further subdivide the newborn OLs into premyelinating and myelinating OLs, we determined that *Lrp1*-deletion significantly increased the proportion that were myelinating OLs (**Fig. 4F**), confirming that *Lrp1*-deletion enhances adult myelination.

Despite the difference in overall cell number, the morphology of the myelinating OLs added to the brain of control and *Lrp1*-deleted mice was equivalent (**Fig. 4G, H**). Our detailed morphological analysis of individual GFP^+^ myelinating OLs in the motor cortex of LE-control and LE-*Lrp1*-deleted mice revealed that neither the average number of internodes elaborated by GFP^+^ myelinating OLs (**Fig. 4I**) or the mean length of internodes elaborated by GFP^+^ myelinating OLs (**Fig. 4J**) was changed by *Lrp1*-deletion. Additionally, the length distribution, for internodes elaborated by newborn myelinating OLs in the motor cortex of LE-control and LE-*Lrp1*-deleted mice, was equivalent (**Fig. 4K**). These data indicate that LRP1 negatively regulates the number of myelinating OLs produced by OPCs in the healthy adult mouse brain but does not influence their final myelinating profile.

### LRP1 does not influence NaV, AMPA receptor, L- or T-Type VGCC, PDGFRα or LRP2 expression by OPCs

LRP1 has the potential to influence a number of signaling pathways that directly or indirectly regulate oligodendrogenesis. The conditional deletion of *Lrp1* from neurons *in vitro* and *in vivo* increases AMPA receptor turnover and reduce expression of the GluA1 subunit of the AMPA receptor (72). Adult OPCs express AMPA receptors (73–75) that enhance the survival of premyelinating oligodendrocytes during development (76), and glutamatergic signaling regulates OPC proliferation, differentiation (74,77) and migration (75). To determine whether LRP1 regulates AMPA receptor signaling in OPCs, we obtained whole cell patch clamp recordings from GFP-labelled OPCs in the motor cortex of P57+30 control (*Lrp1^fl/fl^ :: Pdgfrα-histGFP*) and *Lrp1*-deleted (*Pdgfrα-CreER^TM^ :: Lrp1^fl/fl^* :: *Pdgfrα-histGFP*) mice (**Fig. 5**). OPCs elicit a large inward voltage-gated (sodium) current (INa) in response to a series of voltage-steps (**Fig. 5A**) and we found that INa amplitude was not affected by LRP1 expression (**Fig. 5B**). The capacitance (approximation of cell size; **Fig. 5C**) of OPCs was also unaffected by LRP1 expression. AMPA receptors were subsequently activated by the bath application of 100μm kainate, which evoked a large depolarizing current in control and *Lrp1*-deleted OPCs (**Fig. 5D, E**). The amplitude of the evoked current was equivalent for control and *Lrp1*-deleted OPCs across all voltages examined (**Fig. 5E**), suggesting that *Lrp1*-deletion has no effect on the composition or cell-surface expression of AMPA / kainate receptors.

**Figure 5:**
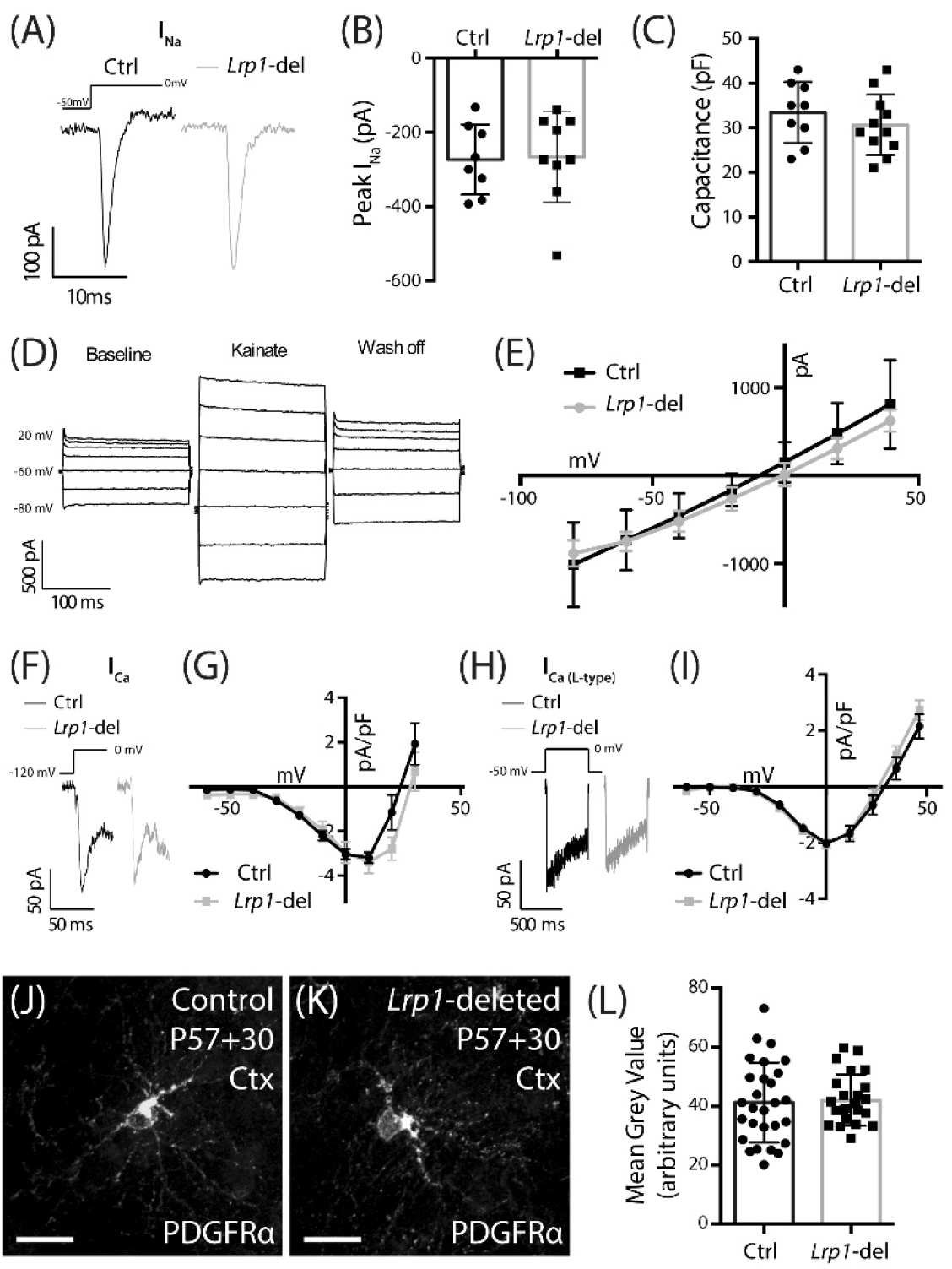
LRP1 does not alter functional NaV, VGCC or AMPA / kainate receptor expression, or total PDGFRα expression in OPCs. (**A**) Representative traces of voltage-gated sodium channels currents evoked in GFP^+^ OPCs in the motor cortex of P57+30 control (*Pdgfrα-histGFP :: Lrp1^fl/fl^*) and *Lrp1*-deleted (*Pdgfrα-CreER™:: Pdgfrα-histGFP :: Lrp1^fl/fl^*) mice. (**B**) Quantification of peak inward voltage-gated sodium current (n≥8 GFP^+^ OPCs analysed from n= 3 mice per genotype; unpaired t-test, p=0.8). (**C**) Quantification of cell capacitance (n≥9 GFP^+^ OPCs analysed from n= 3 mice per genotype; unpaired t-test, p=0.9). (**D**) Representative trace from a control GFP^+^ OPC responding to the bath application of 100μM kainate. (**E**) The current density-voltage relationship of AMPA / kainate receptors in control (n=3 GFP^+^ OPCs) and *Lrp1*-deleted (n=3 GFP^+^ OPCs) cells [mean ± SEM; 2-way repeated measures ANOVA: *Genotype F* (1, 28) = 0.91, p=0.3; *Voltage F* (6, 28) = 31.3, p<0.0001; *Interaction F* (6, 28) = 0.25, p=0.9]. (**F**) Representative traces show the fast inactivating leak subtracted ICa evoked in GFP^+^ OPCs in response to a depolarising step. (**G**) The current density-voltage relationship for the leak subtracted ICa (peak amplitude) recorded from control cells (dark circles, n=11 GFP^+^ OPCs across n=3 mice) and *Lrp1*-deleted cells (grey squares, n=10 GFP^+^ OPCs across n=3 mice) [mean ± SEM; 2-way repeated measures ANOVA: *Genotype F* (1, 190) = 2.85, p=0.09; *Voltage F* (9, 190) = 23.5, p<0.0001; *Interaction F* (9, 190) = 1.14, p = 0.3]. (**H**) Representative traces show the leak subtracted ICa L-type evoked in GFP^+^ OPCs in response to a depolarising step. (**I**) The current density-voltage relationship for leak subtracted ICa L-type (mean sustained current) recorded from control cells (dark circles, n= 7 GFP^+^ OPCs across n=3 mice) and *Lrp1*-deleted cells (grey squares, n=11 GFP^+^ OPCs across n=3 mice) [mean ± SEM; 2-way repeated measures ANOVA: *Genotype F* (1, 176) = 1.03, p=0.3; *Voltage F* (10, 176) = 66.8, p<0.0001; *Interaction F* (10, 176) = 0.62, p=0.8]. (**J-K**) Compressed z-stack confocal image of PDGFRα^+^ OPCs in the motor cortex (Ctx) of P57+30 control and *Lrp1*-deleted mice. (**L**) The mean grey value of PDGFRα staining for individual OPCs measured in the motor cortex of P57+30 control and *Lrp1*-deleted mice (mean ± SD; n ≥ 24 OPCs measured across n=3 mice per genotype; unpaired t-test, p = 0.8). Scale bars represent 17 μm.

LRP1 has also been shown to regulate the cell surface expression and distribution of N-type voltage gated calcium channels (VGCC) by interacting with the α_2_δ subunit (35). In adult OPCs, the closely related L-type VGCCs have been shown to reduce OPC proliferation in the motor cortex and corpus callosum (52), and influence the maturation of OPCs into OLs *in vitro* (78). The other major VGCCs expressed by OPCs are T-type VGCCs (60,79) which are activated at lower (hyperpolarized) voltages than L-type channels and inactivate quickly (transient). To determine whether the distribution of VGCCs is altered following *Lrp1* deletion, we performed whole cell patch clamp electrophysiology and measured the current density (pA/pF) in OPCs from control and *Lrp1*-deleted mice (**Fig. 5F-I**). We found that the VGCC current density was equivalent for OPCs in the motor cortex of control and *Lrp1*-deleted mice (**Fig. 5F, G**). The current density was also equivalent between OPCs from control and *Lrp1*-deleted mice when measured currents were elicited selectively through L-type VGCCs (**Fig. 5H, I**), indicating that LRP1 does not influence L- or T-type VGCC expression in adult OPCs.

Tissue plasminogen activator (tPA) is an LRP1 ligand (80,81), and its addition to astrocytic cultures increases PDGF-CC cleavage and activation (33). While PDGF-CC is a ligand of PDGFRα, a key receptor regulating OPC proliferation, survival and migration 66,67,82), increased mitogenic stimulation would not account for LRP1 reducing OPC proliferation. In other cell types, LRP1 has instead been shown to influence the cell surface expression of PDGFRβ (63,83), a receptor that is closely related to PDGFRα. When performing immunohistochemistry using an antibody against the intracellular domain of PDGFRα, it is not possible to specifically quantify the cell surface expression of PDGFRα in OPCs with and without LRP1, however we were able to quantify PDGFRα expression (mean grey value; **Fig. 5J-l**), and determined that LRP1 did not influence total PDGFRα expression.

The low-density lipoprotein receptor related protein 2 (LRP2) is a large cell surface receptor that is closely related to LRP1, with a number of common ligands (84). LRP2 can increase the proliferation of neural precursor cells in the subependymal zone (85), and the proliferation and survival of skin cancer cells (86), however, it is unclear whether cells of the OL lineage express LRP2 (22–24), or whether *Lrp1*-deletion could alter LRP2 expression. We examined this possibility by performing immunohistochemistry on coronal brain cryosections from P57+30 control and *Lrp1*-deleted mice to detect LRP2 and PDGFRα or ASPA (**Fig. S2**). We determined that LRP2 is not expressed by OPCs or OLs in mice of either genotype, despite the robust expression of LRP2 by Iba1^+^ microglia (**Fig. S2**). These data indicate that compensation from LRP2 or a change in LRP2 expression by OPCs is not responsible for the elevated OPC proliferation and differentiation observed in *Lrp1*-deleted mice.

### LRP1 ligand-mediated activation and *Lrp1*-deletion do not alter OPC proliferation *in vitro*

Our data suggest that in the healthy adult mouse CNS, *Lrp1*-deletion either increases OPC proliferation which then results in an increased number of newborn OLs, or increases OPC differentiation, which subsequently triggers a homeostatic increase in OPC proliferation to maintain the OPC population. Previous studies have shown that *Lrp1* deletion could enhance the proliferation of retinal endothelial cells (87), while the activation of LRP1 by tPA could enhance the proliferation of interstitial fibroblasts (88). To determine whether LRP1 directly suppresses OPC proliferation, we generated primary OPC cultures from the cortex of P0-P5 control (*Pdgfrα-hGFP*) or *Lrp1*-deleted (*Pdgfrα-hGFP :: Lrp1^fl/fl^*) mice. After 7 days *in vitro* (DIV), OPCs were incubated with 1 μM TAT-Cre for 90 min, and LRP1 expression was determined at 9 DIV by performing immunocytochemistry to detect PDGFRα (red), GFP (green) and LRP1 (blue) (**Fig. 6A, B**). Following Tat-Cre treatment all OPCs cultured from control mice expressed LRP1, however, only ~21% of PDGFRα^+^ OPCs cultured from *Lrp1*-deleted mice retained LRP1 expression (**Fig. 6C**). At the same time-point, additional control and *Lrp1*-deleted OPC cultures were exposed to EdU, to label all cells that entered the S-phase of the cell cycle over a 10-hour period. By performing immunocytochemistry to detect GFP (green), LRP1 (red) and EdU (**Fig. 6D, E**), we found that LRP1 expression did not influence OPC proliferation *in vitro*, as the fraction of PDGFRα^+^ OPCs that were EdU^+^ was equivalent in control and *Lrp1*-deleted cultures (**Fig. 6F**).

**Figure 6:**
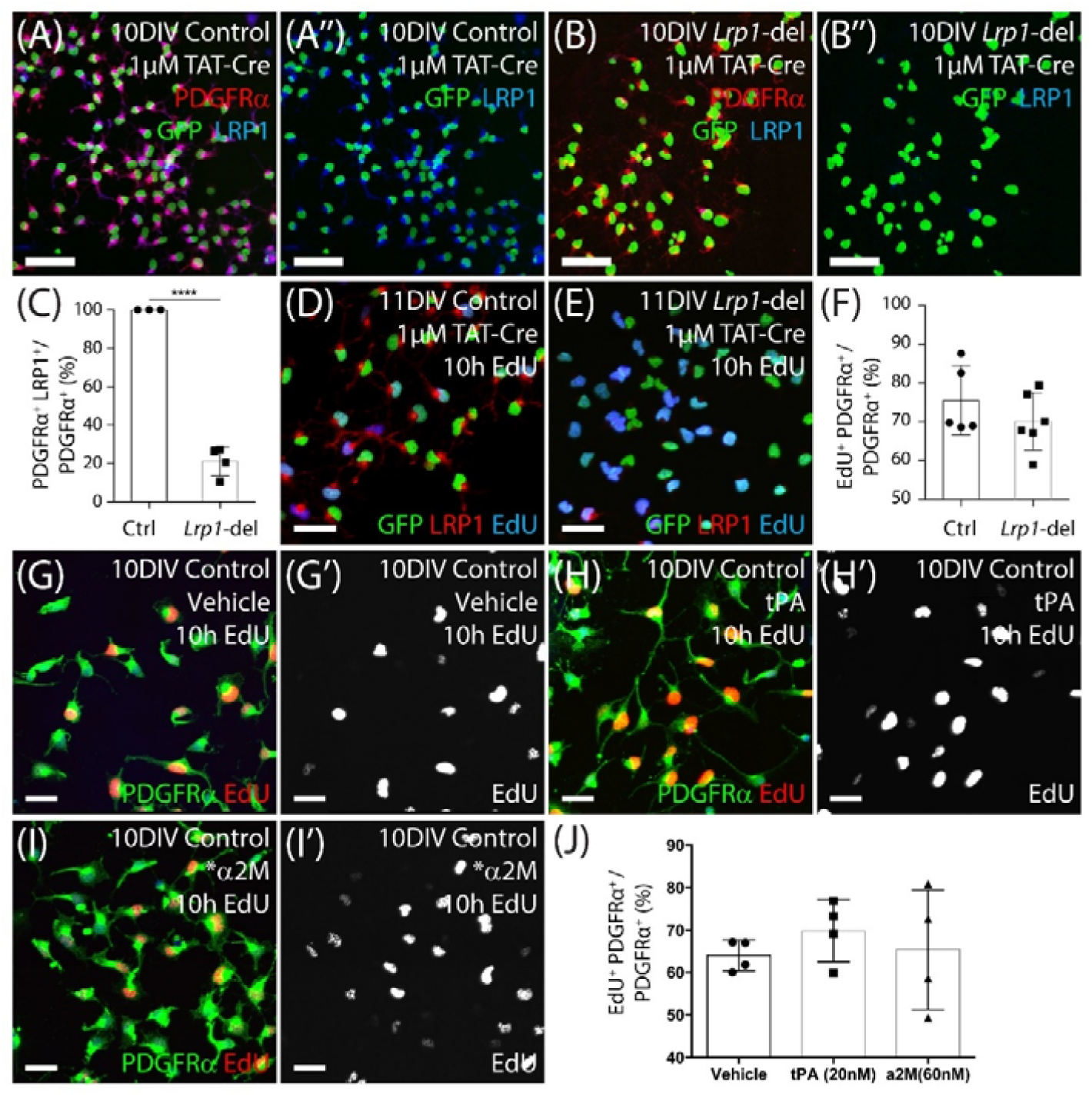
LRP1 does not affect OPC proliferation in vitro. (**A-B**) Tat-Cre-treated OPCs cultured from the cortex of early postnatal control (*Pdgfrα-histGFP*) and *Lrp1*-deleted (*Pdgfrα-histGFP :: Lrp1^fl/fl^*) mice were immunolabelled to detect PDGFRα (red), LRP1 (blue) and GFP (green). (**C**) Quantification of the proportion (%) of control and *Lrp1*-deleted OPCs that express LRP1 48 hours after TAT-Cre treatment (mean ± SEM, n≥3 independent cultures per genotype; unpaired t-test, **** = p<0.0001). (**D-E**) Tat-Cre-treated OPCs from control and *Lrp1*-deleted mice exposed to EdU for 10 hours and immunolabelled to detect GFP (green), LRP1 (red) and EdU (blue). (**F**) Quantification of the proportion (%) of control and *Lrp1*-deleted OPCs that become EdU over a 10-hour labelling period (mean ± SEM, n≥5 independent cultures per genotype; unpaired t-test, p= 0.3). (**G-I**) Compressed confocal z-stack images showing OPCs cultured from control mice that were exposed to EdU and either vehicle (g-g’), 20nM tPa (H-H’) or 60nM *α2M (I-I’) for 10 hours and processed to detect EdU (red) and PDGFRα (green). (**J**) Quantification of the proportion (%) of control OPCs that incorporated EdU when treated with vehicle, tPa or *α2M for 10 hours (mean ± SEM, n≥4 independent cultures; 1-way ANOVA: *Treatment F* (2, 9) = 0.42, p=0.66). Scale bars represent 17 μm. DIV = days *in vitro*; tPA = tissue plasminogen activator; *α2M = activated α-2 macroglobulin.

To further confirm that LRP1 activation by ligands does not directly influence OPC proliferation, we added vehicle (milliQ water) or the LRP1 ligands tPA (20nM) or activated α-2 macroglobulin (*α2M; 60mM) to OPC primary cultures for 10 hours, along with EdU (**Fig. 6G-J**). By performing immunocytochemistry to detect PDGFRα (green) and EdU (red), we determined that the proportion of OPCs that became EdU-labelled did not change with the addition of tPA or *α2M (**Fig 6J**), indicating that LRP1 activation by these ligands is unable to modify OPC proliferation *in vitro*. These data suggest that LRP1 does not have a direct or cell intrinsic effect on OPC proliferation.

### *Lrp1*-deletion increases OPC differentiation *in vitro* but the ligand activation of LRP1 does not

*In vitro*, OPCs can be triggered to differentiate by withdrawing the mitogen PDGF-AA and providing triiodothyronine (T3) in the culture medium. To determine whether *Lrp1* deletion can enhance OPC differentiation, Tat-Cre-treated control and *Lrp1*-deleted OPCs were transferred into differentiation medium for 4 days before they were immuno-labelled to detect PDGFRα^+^ OPCs (red) and MBP^+^ OLs (green) (**Fig. 7A, B**). We determined that the proportion of cells that were PDGFRα^+^ OPCs was reduced in the *Lrp1*-deleted cultures, while the proportion of cells that were MBP^+^ OLs was significantly increased compared with control cultures (**Fig. 7C**).

**Figure 7:**
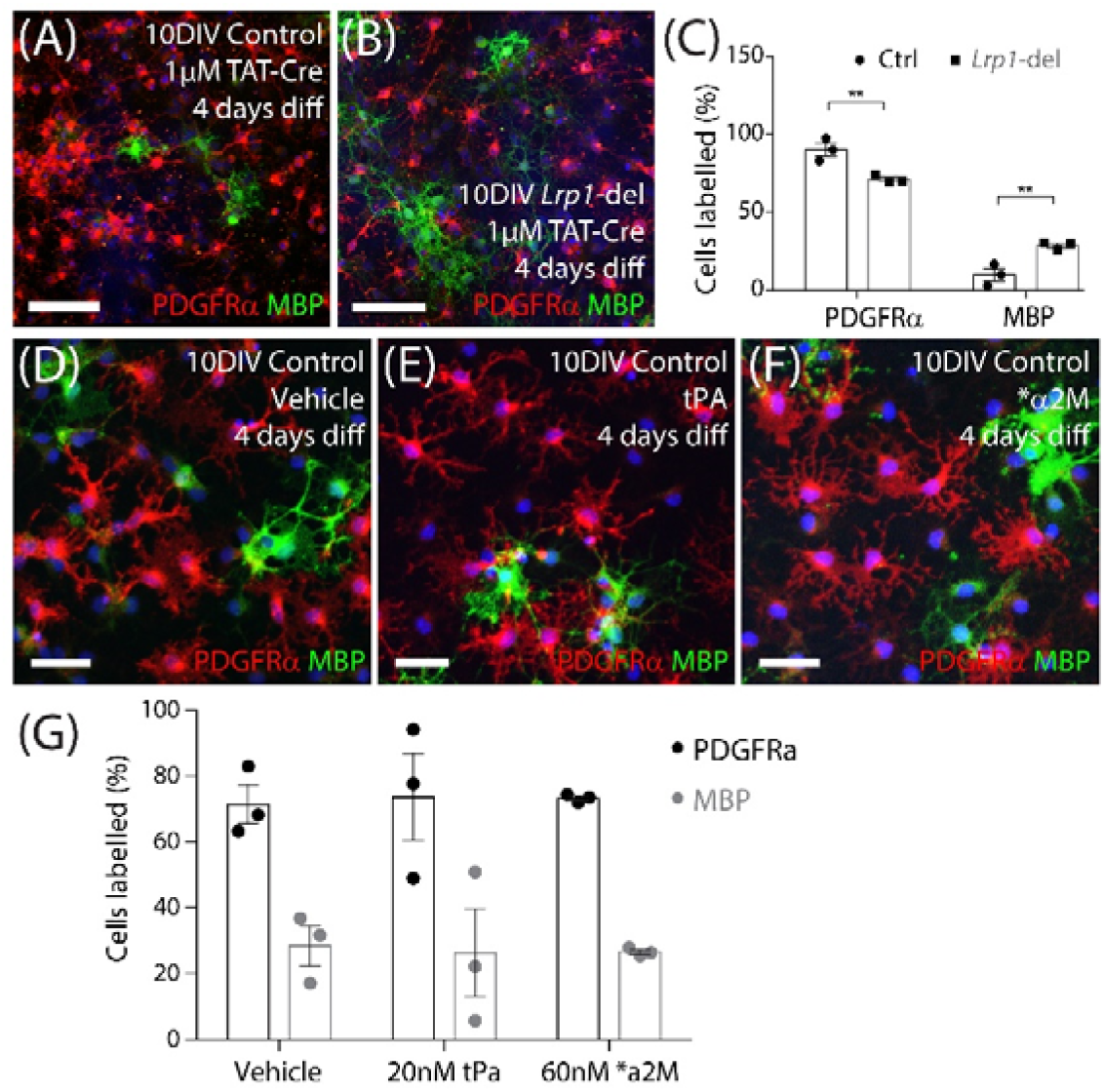
LRP1 expression reduces the OPC differentiation *in vitro*. (**A-B**) Compressed confocal z-stack images of differentiated control and Lrp1-deleted (Lrp1^fl/fl^) OPC cultures immunolabelled to detect OPCs (PDGFRα, red), OLs (MBP, green) and all cell nuclei (Hoescht 33342, blue). OPCs were exposed to Tat-Cre, cultured for a further 48-hours, and then transferred to differentiation medium for 4 days. (C) Quantification of the proportion (%) of cells that were PDGFRα^+^ OPCs or MBP^+^ OLs in control and Lrp1-deleted cultures [mean ± SEM for n=3 independent cultures per genotype; 2-way ANOVA: Cell type F (1, 8) = 212, p <0.0001; Genotype F (1, 8) = 0.00006, p= 0.97; Interaction F (1, 8) = 19.9, p=0.02]. Bonferroni posthoc test ** p = 0.005. (D-F) Compressed confocal z-stack images of OPCs from control mice that transferred into differentiation medium and exposed to vehicle (DMSO; D), tPa (E) or *α2M (F) for 4 days before being immunolabelled to detect OPCs (PDGFRα, red), OLs (MBP, green) and all cell nuclei (Hoescht 33342, blue). (G) Quantification of the proportion (%) of OPCs that became PDGFRα^+^ OPCs or MBP^+^ OLs after 4 days in differentiation medium with vehicle, tPa or *α2M (mean ± SEM, n=3 independent cultures per treatment; 2-way ANOVA: Cell type F (1, 12) = 44.6, p<0.0001; Treatment F (2, 10) = 2.57e-012, p>0.99; Interaction F (2, 12) = 0.03, p= 0.97]. tPa = tissue plasminogen activator; *α2M = activated α2 macroglobulin. Scale bars represent 34 μm (A, B) or 17 μm (D-F).

To determine whether the ligand activation of LRP1 was sufficient to suppress OPC differentiation, OPC primary cultures were instead transferred into differentiation medium containing vehicle, tPA (20nM) or *α2M (60nM) for 4 days. By performing immunocytochemistry to detect PDGFRα^+^ OPCs and MBP^+^ OLs (**Fig. 7D-F**) we found that the activation of LRP1 by tPA or *α2M had no impact on the proportion of cells that differentiated over time (**Fig. 7G**). These data suggest that LRP1 normally acts to suppress OPC differentiation, however, this effect is independent of tPA and α2M signaling. Furthermore, the effect of LRP1 on OPC proliferation *in vivo* is likely to be a secondary consequence of the influence that LRP1 exerts on OPC differentiation.

### OPC specific *Lrp1* deletion reduced lesion volume in the cuprizone mouse model of demyelination

Having shown that *Lrp1* deletion increases adult OPC differentiation and consequently myelination, we wanted to determine whether the deletion of *Lrp1* from OPCs could improve remyelination.

Control (*Pdgfrα-CreER^TM^ :: Rosa26-YFP*) and *Lrp1*-deleted (*PdgfrαCreER^TM^ :: Rosa26-YFP :: Lrp1^fl/fl^*) mice received tamoxifen by oral gavage at P57, and at P64 were transferred onto a diet containing 0.2% (w/w) cuprizone. Cuprizone feeding induces significant OL loss and demyelination of the corpus callosum, but also triggers oligodendrogenesis. After 5 weeks of cuprizone feeding, control and *Lrp1*-deleted mice were perfusion fixed and coronal brain sections stained to detect myelin by black-gold staining (**Fig. 8**). We detected overt demyelination in the corpus callosum of control and *Lrp1*-deleted mice (**Fig. 8A-D**), however, *Lrp1*-deleted mice had significantly less demyelination than controls (**Fig. 8E**).

**Figure 8:**
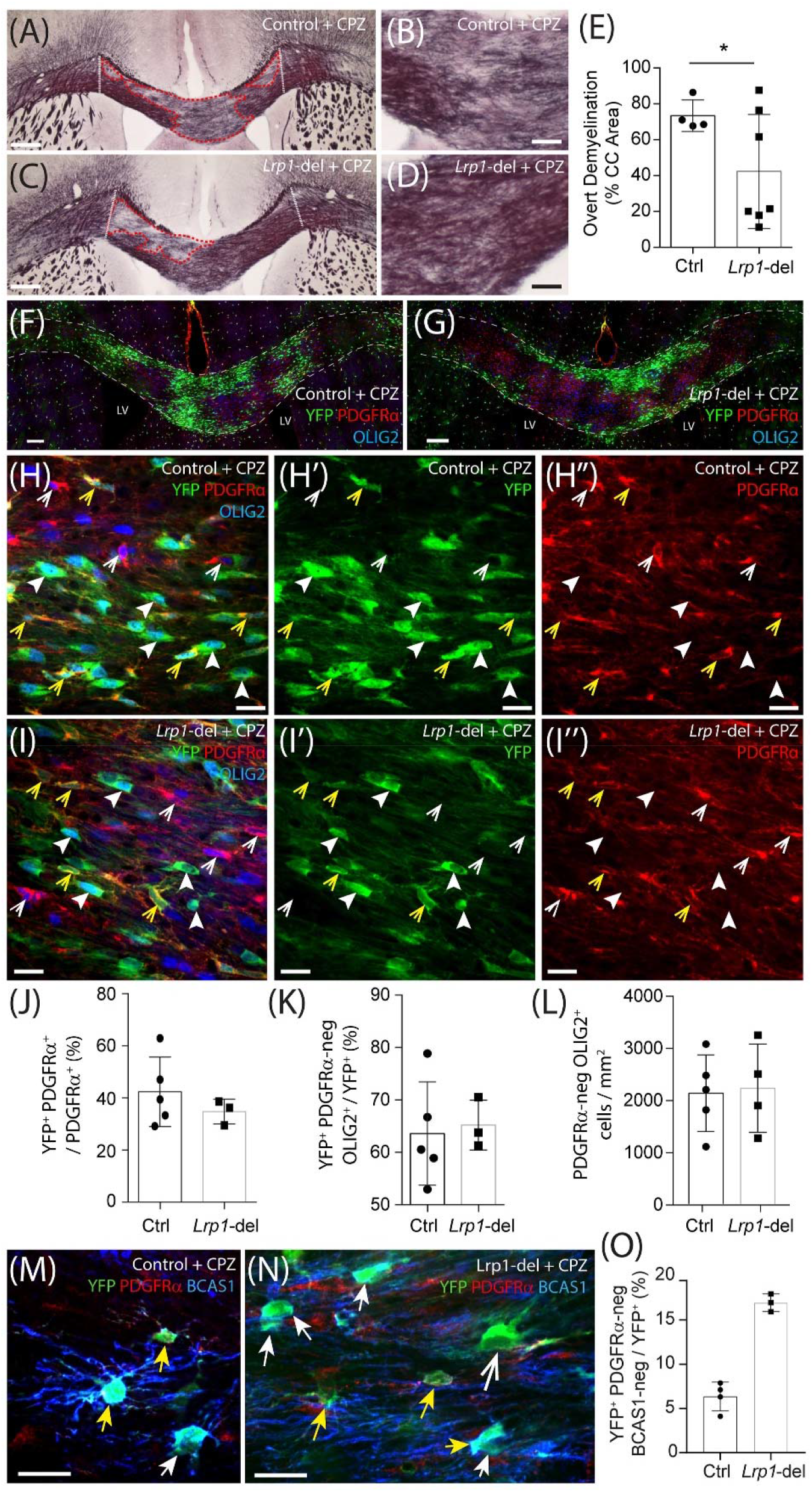
Lrp1-deletion enhances callosal remyelination and oligodendrocyte maturation. (**A-D**) Young adult control (*Pdgfrα-CreER^TM^ :: Rosa26-YFP*) and *Lrp1*-deleted (*Pdgfrα-CreER^TM^ :: Rosa26-YFP :: Lrp1^fl/fl^*) mice were fed a diet containing 0.2% cuprizone for 5 weeks. At the end of 5 weeks, coronal brain sections were collected and stained with black-gold to visualise myelin in the corpus callosum. Dotted white lines define the lateral limits of the region of the corpus callosum analysed. Red dashed lines denote areas of overt demyelination. (**E**) The proportion (% area) of the corpus callosum with feint or absent black-gold staining (severe demyelination) in cuprizone-fed control and *Lrp1*-deleted mice (mean ± SD, n ≥ 4 mice per genotype; unpaired t-test with Welch’s correction, * p=0.04). (**F-I**) Compressed confocal z-stack images of the corpus callosum in cuprizone-fed control and *Lrp1*-deleted mice immunolabelled to detect YFP (green), PDGFRα (red) and OLIG2 (blue). White dashed lines indicate the boundary of the white matter tract; yellow arrowheads indicate YFP^+^ PDGFRα^+^ parenchymal OPCs; solid white arrowheads indicate YFP^+^ PDGFRα-negative newborn OLs; white arrowheads indicate YFP-negative PDGFRα^+^ stem cell-derived OPCs. (**J**) Quantification of the proportion (%) of OPCs in the corpus callosum of cuprizone-fed control and *Lrp1*-deleted mice that are YFP^+^, PDGFRα^+^ parenchymal OPCs (mean ± SD, n ≥ 3 mice per genotype; unpaired t-test, p=0.4). (**K**) Quantification of the proportion of YFP^+^ cells in the corpus callosum of cuprizone-fed control and *Lrp1*-deleted mice that are PDGFRα-negative OLIG2^+^ newborn OLs (mean ± SD, n ≥ 3 mice per genotype; unpaired t-test, p=0.9). (**L**) Quantification of the density of OLIG2^+^ OLs in the corpus callosum of cuprizone-fed control and *Lrp1*-deleted mice (mean ± SD, n ≥ 4 mice per genotype; unpaired t-test, p=0.9). (**M-N**) Single z-plane confocal images from the corpus callosum of cuprizone-fed control and *Lrp1*-deleted mice stained to detect YFP (green) PDGFRa (red) and BCAS1 (blue). Solid yellow arrows indicate parenchymal OPCs (YFP^+^ PDGFRa^+^ ± BCAS1). Solid white arrows indicate newborn premyelinating OLs derived from parenchymal OPCs (YFP^+^ PDGFRa-neg BCAS1^+^). Large white arrow indicates newborn mature OL derived from parenchymal OPCs (YFP^+^ PDGFRa-neg BCAS1-neg). (**O**) Quantification of the proportion of YFP+ cells in the corpus callosum of cuprizone-fed control and *Lrp1*-deleted mice that are newborn mature OLs (YFP^+^ PDGFRa-neg BCAS1-neg). Scale bars represent 150 μm (A, C), 30 μm (B, D), 100 μm (F, G), 17 μm (H, I) or 20 μm (M, N). LV= lateral ventricle.

When mice received EdU via their drinking water from week 2 to week 5 of cuprizone feeding, we found that the vast majority of OLIG2^+^ cells in the corpus callosum of control and *Lrp1*-deleted mice became EdU-labelled during this period (**Fig S3**). Despite being newborn cells, many of the PDGFRα^+^ OPCs (red) and PDGFRα-negative OLIG2^+^ OLs (blue) within the corpus callosum of control and *Lrp1*-deleted mice did not co-label with YFP (green) (**Fig. 8F, G**), indicating that these cells were not derived from the YFP^+^ parenchymal OPC population. Following cuprizone-induced demyelination, both parenchymal OPCs (YFP-labelled) and neural stem cell-derived OPCs (YFP-negative) contribute to OL replacement and remyelination (89). As the proportion of OPCs that were YFP^+^ parenchymal OPCs was equivalent in the corpus callosum of control and *Lrp1*-deleted mice after cuprizone demyelination (**Fig 8J**), and total OPC density was unaffected by genotype (838 ± 165 OPCs / mm^2^ in control and 740 ± 134 OPCs / mm^2^ in *Lrp1*-deleted corpus callosum; mean ± SD for n = 5 control and n=3 *Lrp1*-deleted mice; unpaired t-test, p = 0.67), we can conclude that the expression of LRP1 by parenchymal OPCs does not influence OPC production by neural stem cells.

Following demyelination, YFP^+^ parenchymal OPCs present in the corpus callosum of *Lrp1*-deleted mice lacked LRP1, however, the YFP-negative neural stem cell-derived OPCs had intact LRP1 expression (**Fig. S3**). Furthermore, parenchymal OPCs no longer generated more OLs in *Lrp1*-deleted mice compared to controls, as 60% ± 15% of YFP^+^ cells were PDGFRα-negative OLIG2^+^ newborn OLs in the corpus callosum of control mice and 65% ± 5% of YFP^+^ cells were PDGFRα-negative, OLIG2^+^ newborn OLs in the corpus callosum of *Lrp1*-deleted mice (**Fig. 8K**; mean ± SD for n=5 control and n=3 *Lrp1*-deleted mice). As total OL density was also equivalent in the corpus callosum of control and *Lrp1*-deleted mice (**Fig. 8L**), a change in oligodendrogenesis could not account for the reduced lesion size detected in *Lrp1*-deleted mice. However, by performing immunohistochemistry to detect YFP, the OPC marker PDGFRα, and Breast Carcinoma Amplified Sequence 1 (BCAS1), a protein expressed by some OPCs and all pre-myelinating OLs (90,91), we were able to determine that the fraction of YFP^+^ cells that were mature OLs (YFP^+^ PDGFRα-neg BCAS1-neg) was increased in the corpus callosum of *Lrp1*-deleted mice compared to controls (**Fig. 8M-O**). These data suggest that adult OPCs express LRP1 that acts to suppress the production of mature, myelinating OLs in the healthy and injured CNS of adult mice.

## Discussion

Within the OL lineage, LRP1 is highly expressed by OPCs and rapidly down-regulated upon differentiation (22,23,25,50), suggesting that LRP1 regulates the function or behavior of the progenitor cells. As LRP1 can signal in a number of different ways (26,27,92), and has been shown to influence cellular behaviours relevant to OPCs, such as proliferation, differentiation (48,87,93) and migration (94–98), we took a conditional gene deletion approach to determine whether *Lrp1* influenced the behavior of adult mouse OPCs. We report that LRP1 is a negative regulator of adult oligodendrogenesis in the healthy adult mouse CNS. *Lrp1*-deletion increased the number of OPCs that differentiated into OLs, including the production of mature, myelinating OLs. However, *Lrp1*-deletion did not alter the number or length of internodes produced by the myelinating OLs. Following cuprizone-induced demyelination, when the drive for oligodendrogenesis is increased but callosal OPCs are exposed to myelin debris and an environment that promotes glial activation, we found that LRP1 no longer influenced the number of newborn OLs added to the corpus callosum, but did impair the maturation of the newborn OLs.

### Why does *Lrp1*-deletion have a delayed effect on OPC proliferation in the healthy adult mouse CNS?

At any one time, the majority of OPCs in the healthy adult mouse CNS are in the G0 phase of the cell cycle (62). In young adulthood, all OPCs in the corpus callosum re-enter the cell cycle and divide at least once in a 10-day period, but a similar level of turnover takes ~38 days for OPCs in the cortex (4). In this study, we found that *Lrp1*-deletion increased the rate at which OPCs re-entered the cell cycle, but the onset of this phenotype was not coincident with *Lrp1*-deletion. More specifically, 7 days after tamoxifen delivery, *Lrp1*-deletion did not alter the rate at which OPCs entered S-phase of the cell cycle, as an equivalent proportion of the OPC population became EdU labelled over time in control and *Lrp1*-deleted mice (**Fig. 2**). However, when the analysis was delayed by another 25 days (32 days after tamoxifen), we found that the rate of EdU labelling was significantly higher for OPCs in the corpus callosum of *Lrp1*-deleted mice relative to controls. It is feasible that LRP1 directly suppresses OPC proliferation, as LRP1 is known to modulate the proliferation of other cell types (48,87,93,99–101), suppressing the hypoxia-induced proliferation of mouse and human retinal endothelial cells by regulating the activity of poly (ADP-ribose) polymerase-1 (PARP-1) (87), and suppressing the proliferation of cultured mouse vascular smooth muscle cells by reducing PDGFRβ activity (101,102). However, the inability of *Lrp1*-deletion to acutely influence OPC proliferation *in vivo*, or directly influence OPC proliferation *in vitro* [**Fig. 6**; (49)], suggests that LRP1 indirectly affects OPC proliferation.

OPC proliferation is intimately linked to OPC differentiation *in vivo* (6). As the number of new OLs that are added to the adult mouse brain increases following the conditional deletion of *Lrp1* from adult OPCs (**Fig. 3** and **Fig. 4**), it is possible that LRP1 indirectly suppresses OPC proliferation by directly suppressing OPC differentiation. This initially seemed unlikely as a previous report indicated that *Myrf*, *Mbp* and *CNPase* mRNA expression was equivalent in *Olig1-Cre* and *Olig1-Cre :: Lrp1^fl/fl^* OPC cultures after only 2 days of differentiation (50), however, we found that the deletion of *Lrp1* from cultured mouse OPCs was sufficient to increase their differentiation into MBP^+^ OLs over a 4-day period (**Fig. 7**). The direct suppression of OPC differentiation by LRP1 could certainly explain both the increased number of newborn OLs and the increased OPC proliferation detected in the brain of *Lrp1*-deleted mice (see **Fig. 9**), as increased OPC differentiation would stimulate the proliferation of adjacent OPCs, ensuring the homeostatic maintenance of the progenitor pool (6).

**Figure 9:**
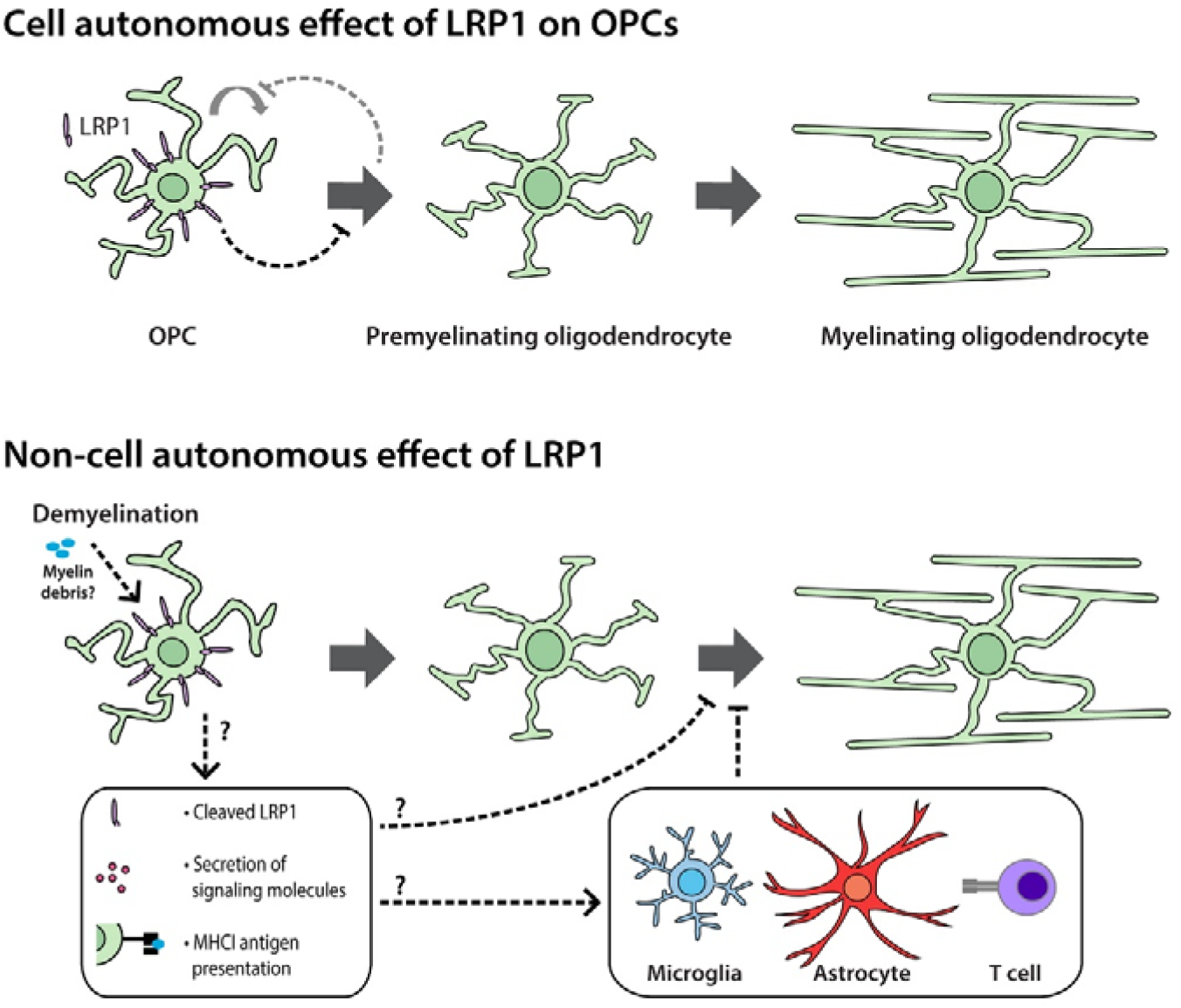
LRP1 signaling in OPCs may influence myelination by different mechanisms in the healthy and demyelinated CNS. Cell autonomous: *In vivo*, LRP1 signaling by adult mouse OPCs reduces OPC proliferation and the number of newborn OLs added to the brain. As LRP1 suppresses OPC differentiation *in vitro*, we propose that LRP1 has a direct, cell autonomous effect on OPCs, primarily suppressing OPC differentiation and having a secondary, homeostatic effect on OPC proliferation. The increased production of new OLs was accompanied by an increase in the addition of new myelinating OLs to the brain. Non cell autonomous: Following cuprizone-induced demyelination, LRP1 no longer suppresses OPC differentiation. This may be because the demyelinating injury alters the nature of LRP1 signaling or pro-oligodendrogenic signals become dominant. However, LRP1 expression by OPCs hinders OL maturation and remyelination. As OLs rapidly lose LRP1 expression during differentiation, LRP1 signaling from OPCs must exert an indirect effect on OL maturation in the remyelinating environment. This may be the result of soluble LRP1 or secreted proteins acting on pre-myelinating OLs or LRP1 initiating the secretion of inflammatory molecules and enabling antigen cross-presentation to influence the behavior of other cells, such as microglia, astrocytes or lymphocytes to influence the maturation of remyelinating OLs.

### LRP1 is a negative regulator of adult oligodendrogenesis

New OLs are added to the adult mouse CNS throughout life (2,3,103,104), however, when we followed the fate of adult OPCs after *Lrp1* deletion, we observed a significant increase in the number of new OLs added to the corpus callosum and motor cortex within 30 and 45 days of gene deletion. By contrast, in the developing mouse optic nerve, deleting *Lrp1* from cells of the OL lineage (*Olig2-Cre :: Lrp1^fl/fl^*) reduced the number of OLs produced and resulted in hypomyelination by P21 (49). This phenotype was largely attributed to the ability of LRP1 to promote cholesterol homeostasis and peroxisome function, and consequently developmental OPC differentiation (49). Differences in the developing and adult brain environments (51), regional differences in signaling between the brain and optic nerve, or changes in gene expression between developmental and adult OPCs (53) could account for this clear difference in LRP1 function. As LRP1 can suppress the differentiation of OPCs cultured from the developing mouse cortex, it seems more likely that LRP1 signaling differs between OPCs in the optic nerve and brain. However, it could also be explained by *Olig2* expression in neural stem / progenitor cells or its transient expression by astrocytes (105,106) resulting in, for example, the unintended deletion of *Lrp1* from some neural stem cells, which is known to reduce the overall generation of cells of the OL lineage (47,48), and would be predicted to impair myelination.

It is important to note that deleting *Lrp1* from adult OPCs not only increased oligodendrogenesis but increased adult myelination. In the healthy adult mouse brain, there is a significant population of pre-myelinating OLs (90,107) that are constantly turned over, as ~78% of newly generated pre-myelinating OLs survive for less than 2 days (9). By using LE-*Pdgfrα-CreER^T2^ :: Tau-mGFP* transgenic mice to visualize the full morphology of the newly generated OLs, we were able to confirm that *Lrp1*-deletion effectively increased the number of newborn, myelinating OLs added to the brain (**Fig. 4**), which equated to a larger number of new myelin internodes being added. However, *Lrp1*-deletion did not seem to directly influence OL maturation in the healthy mouse brain, as the proportion of newborn OLs that were at the pre-myelinating and myelinating stages of differentiation was equivalent between control and *Lrp1*-deleted mice. Furthermore, the myelinating profile of individual myelinating OLs was unaffected by LRP1 expression, as OLs in the cortex of control and *Lrp1*-deleted mice supported the same amount of myelin in an equivalent configuration (**Fig. 4**). Therefore, LRP1 appears to regulate the overall number of new OLs generated in the adult mouse CNS, but not their maturation.

### LRP1 indirectly suppresses callosal remyelination

As *Lrp1*-deletion increased OPC differentiation in healthy adult mice, we predicted that *Lrp1*-deletion would enhance oligodendrogenesis in response to a cuprizone-demyelinating injury. We instead found that OL production was unaffected by LRP1 expression. It has been reported that within 3.5 days of cuprizone withdrawal, *Olig1-Cre :: Lrp1^fl/fl^* mice have more OLs and increased MBP coverage of the corpus callosum than *Olig1-Cre* control mice (50). In our study, it is possible that *Lrp1*-deletion was less able to direct parenchymal OPC differentiation, as a significant number of OPCs within the environment retained LRP expression i.e. the injured corpus callosum contained a mixture of *Lrp1*-deleted OPCs and neural stem cell-derived *Lrp1* replete OPCs. However, this seems unlikely, as a similarly mixed population of LRP1^+^ and LRP1-negative OPCs was present in the cortex of healthy adult LE-*Lrp1*-deleted mice, due to their low recombination efficiency, and yet OPCs continued to produce a larger number of new OLs in the cortex of LE-*Lrp1*-deleted mice when compared with controls. An alternative, and perhaps more likely explanation, is that the cuprizone-induced demyelination acted as a robust stimulus for OPC differentiation (89,108), and effectively masked the effect of LRP1 on oligodendrogenesis.

Despite our observation that parenchymal OPCs produced a similar number of newborn YFP^+^ callosal OLs in control and *Lrp1*-deleted mice, and OL density was also equivalent, we determined that *Lrp1*-deleted mice had significantly more callosal myelin and a greater proportion of the YFP^+^ cells had become mature OLs. This effect is unlikely to be a cell autonomous effect of LRP1, as LRP1 is not expressed by newly generated OLs (25). However, LRP1 signaling may allow OPCs to reduce the maturation of nearby OLs within the injury environment (see **Fig. 9**). Neuroinflammation impairs OL generation (109), and OPCs can modulate neuroinflammation, releasing cytokines in response to interleukin 17 receptor signaling (110), and expressing genes associated with antigen processing and presentation (111,112). LRP1 can bind and phagocytose myelin debris (113–115) and LRP1 expression by OPCs can influence the inflammatory nature of the remyelinating environment. RNA profiling of the remyelinating corpus callosum of *Olig1-Cre* and *Olig1-Cre :: Lrp1^fl/fl^* mice, 3.5 days after cuprizone withdrawal, revealed that inflammatory gene expression was reduced in the *Olig1-Cre :: Lrp1^fl/fl^* mice (50). LRP1 signaling could lead to OPCs secreting pro-inflammatory factors or releasing a cleaved, soluble form of LRP1 to enhance the inflammatory response of nearby microglia (42,43). However, LRP1 may also facilitate antigen presentation by OPCs, as the deletion of *Lrp1* from OPCs reduces their expression of MHC class I antigen presenting genes in the corpus callosum of cuprizone-demyelinated mice, and reduces the ability of OPCs to cross-present antigens to lymphocytes *in vitro* (50). As LRP1 signaling may differentially influence OPC function in the healthy and demyelinated CNS, further research is required to fully elucidate its direct and indirect affect on myelination and remyelination.

## Author Contributions

LA, KMY, LF and BVT developed the project and wrote the manuscript. LA, CLC, REP and KAP carried out the experiments. KMY and LF obtained the funding. LA, CLC, KAP and KMY performed the statistical analyses and generated the figures. KMY, LF and BVT provided supervision.

## Acknowledgments

We thank our colleagues at the University of Tasmania for providing animal services support or providing constructive feedback and suggestions.

## Data Availability Statement

All individual data points are provided in the data figures or in the supplementary data of the manuscript. Requests for any other data files should be directed to the corresponding author.

## Graphical abstract

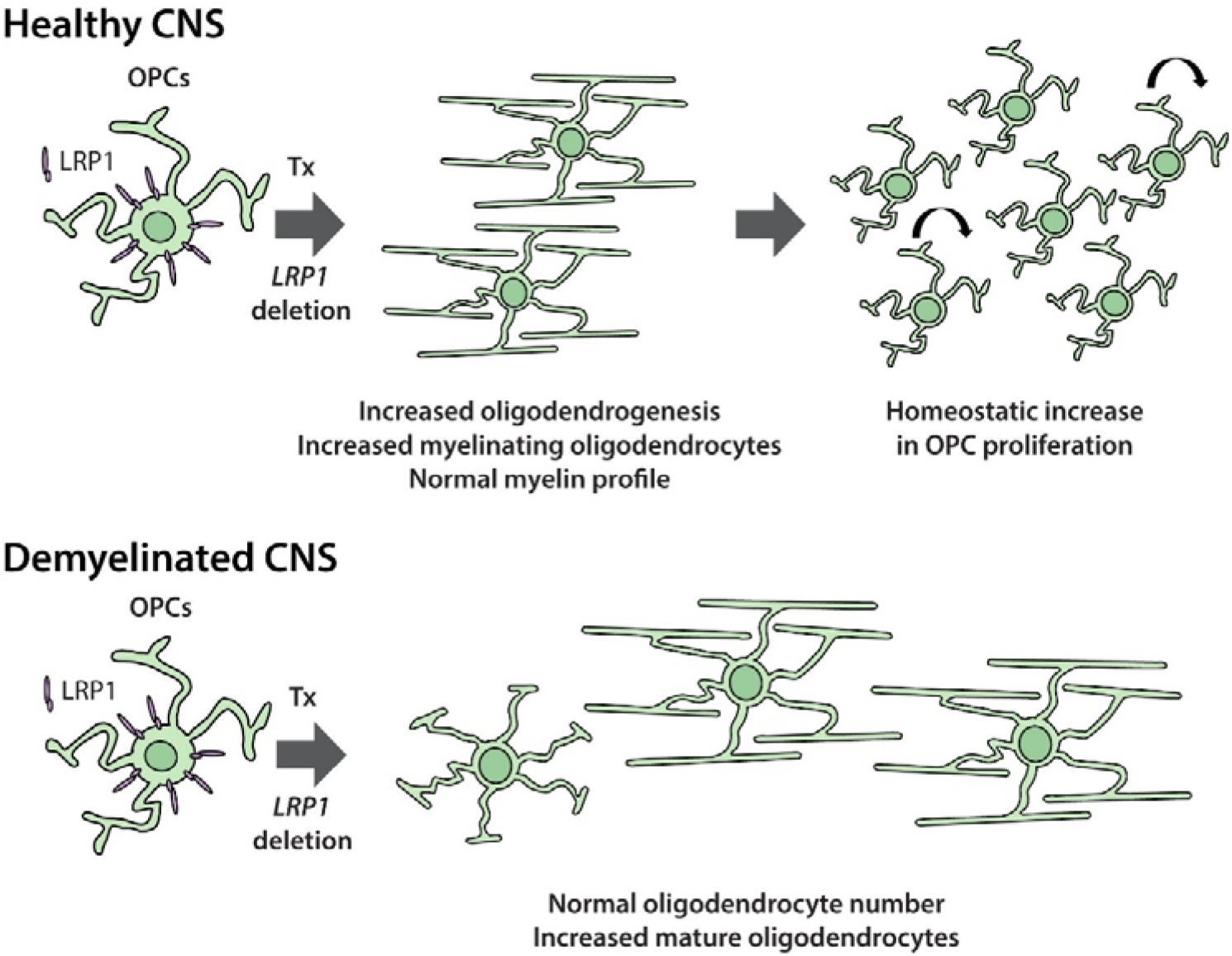

## Supplementary Material – Auderset et al

**Supplementary Figure 1:**
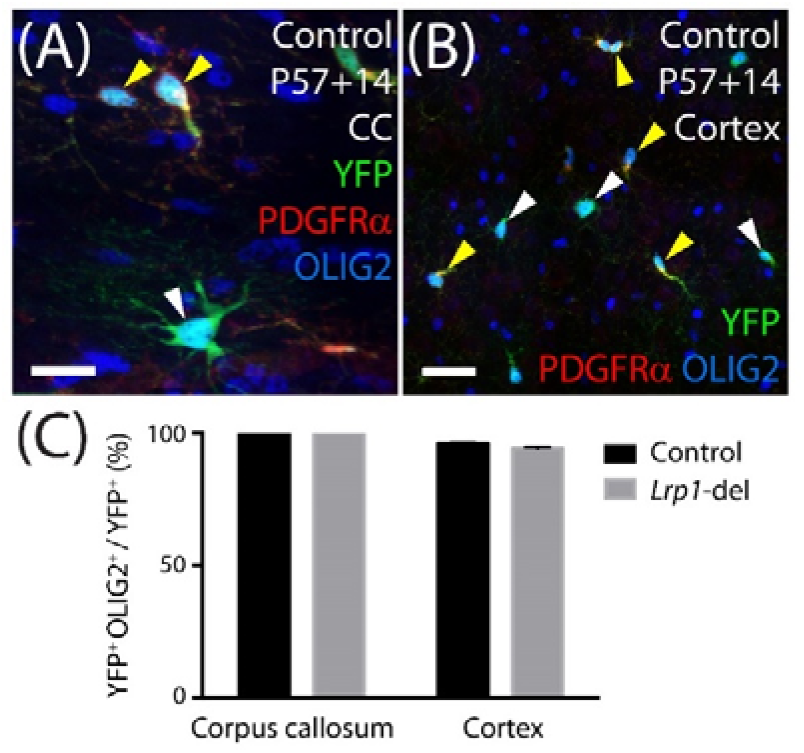
Essentially all YFP-labelled cells belong to the OL lineage. (**A-B**) Confocal images from the corpus callosum (CC) and motor cortex (Cortex) of P57+14 control (*Pdgfrα-CreER^TM^ :: Rosa26YFP*) mice immunolabelled to detect OPCs (PDGFRα, red), YFP (green) and the transcription factor OLIG2 (blue). Solid yellow arrowheads indicate YFP^+^ OLIG2^+^ PDGFRα^+^ OPCs. Solid white arrowheads indicate YFP^+^ OLIG2^+^ PDGFRα-neg newborn OLs. (**C**) Quantification of the proportion (%) of YFP^+^ cells that express OLIG2 in the corpus callosum (100% ± 0% for control and 100% ± 0% for *Lrp1*-deleted) and motor cortex (96.1% ± 0.9% from control and 94.3% ± 1% for *Lrp1*-deleted mice) of P57+14 mice (mean ± SD, n= 3 mice per group). Scale bars represent: 34μm (A) or 17μm (B).

**Supplementary Figure 2:**
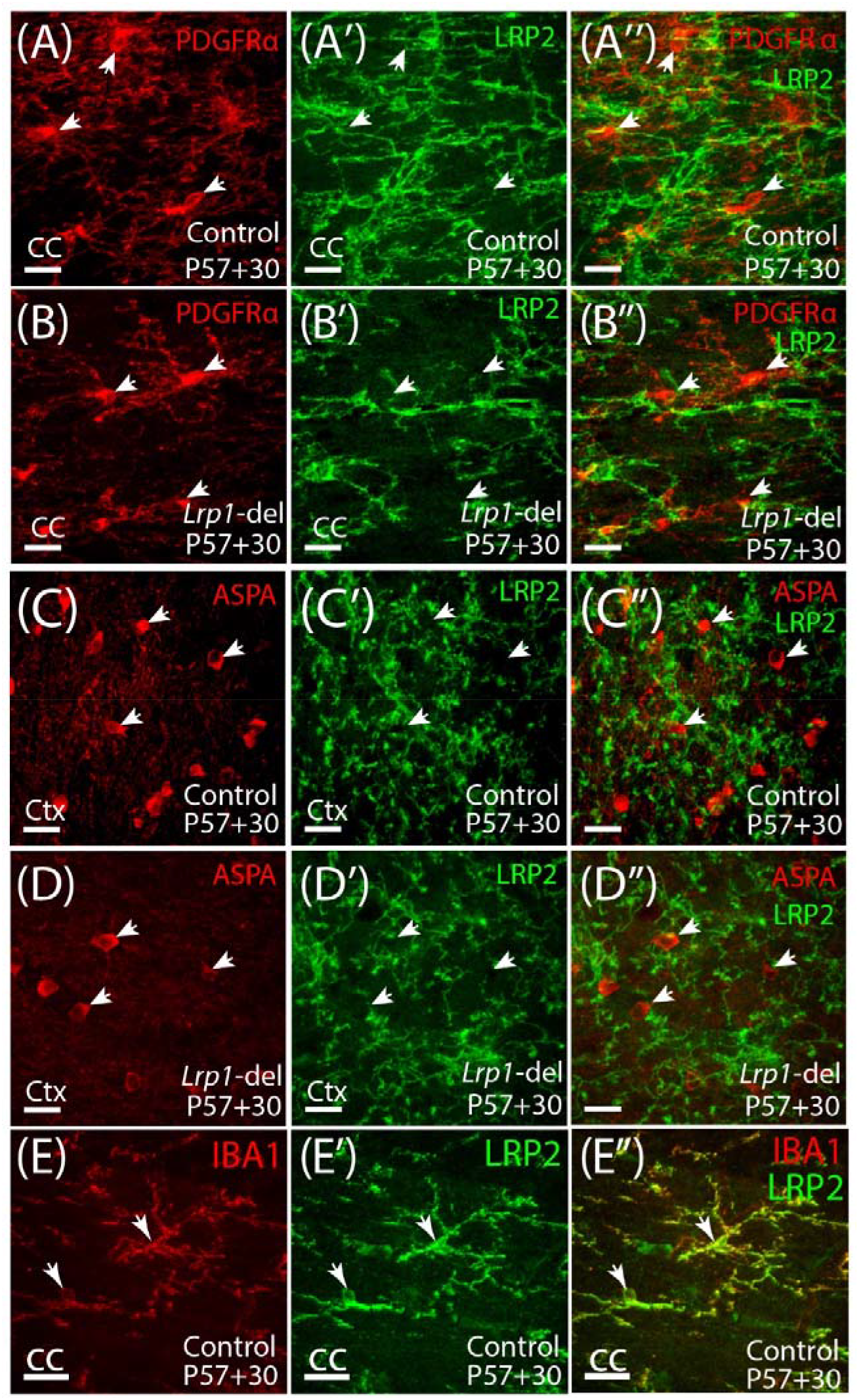
LRP2 is not expressed by OPCs or OLs, but is expressed by microglia. (**A-B**) Compressed z-stack confocal images of the corpus callosum (CC) in a P57+30 control (*Pdgfrα-CreER^TM^*) and *Lrp1*-deleted (*Pdgfrα-CreER^TM^ :: Lrp1^fl/fl^*) mouse, immunolabelled to detect the OPC marker PDGFRα (red) and LRP2 (green). (**C-D**) Compressed z-stack confocal images of the motor cortex (Ctx) in a P57+30 control and *Lrp1*-deleted mouse, immunolabelled to detect the OL marker ASPA (red) and LRP2 (green). (**E**) Compressed z-stack confocal image of the CC in a P57+30 control mouse immunolabelled to detect the microglial marker IBA1 (green) and LRP2 (red). White arrows denote the location of the OPCs (A, B), OLs (C, D) or microglia (E). Scale bars represent 17μm.

**Supplementary Figure 3:**
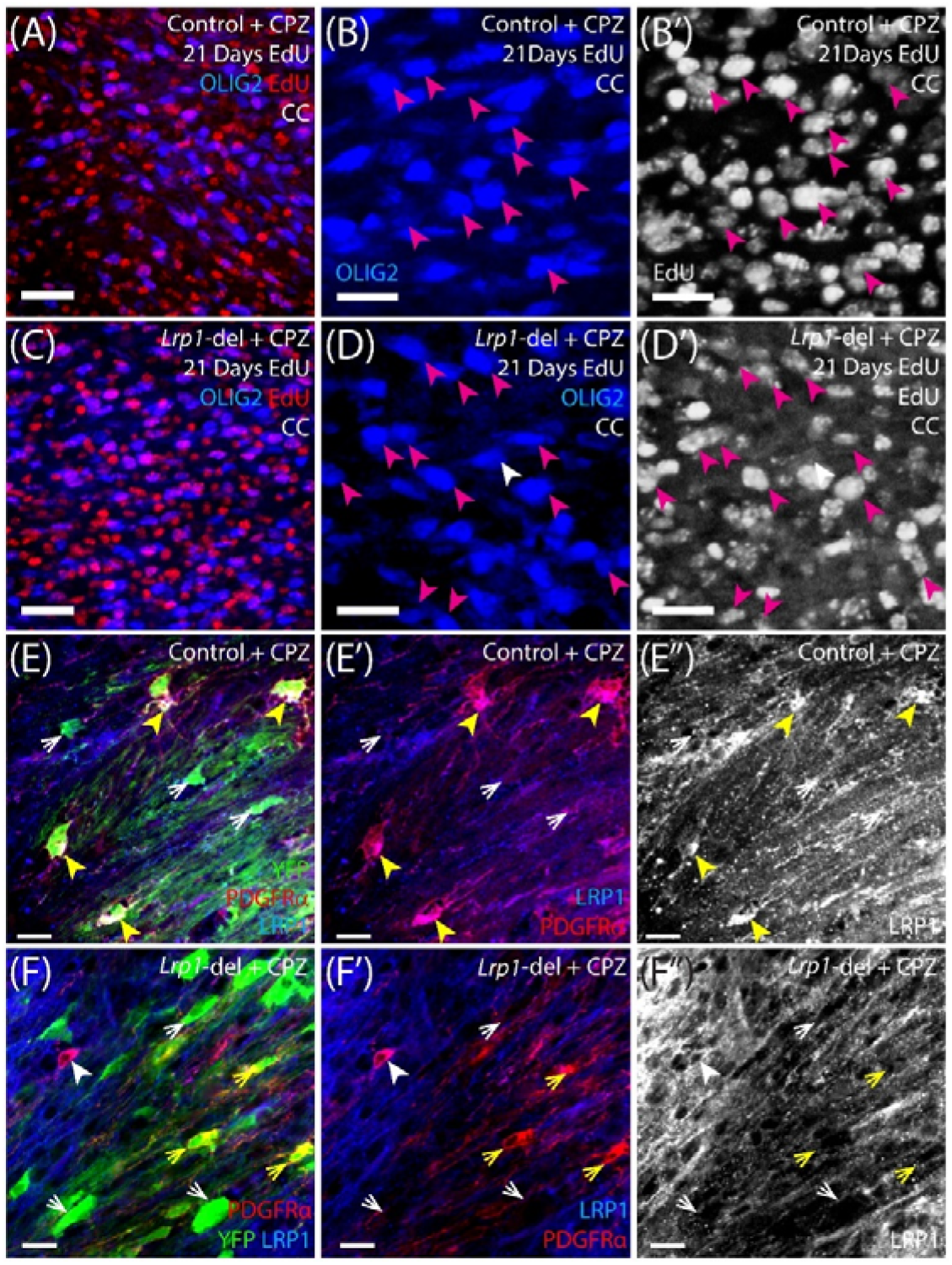
The vast majority of OLIG2^+^ cells present in the corpus callosum of cuprizone-fed control and *Lrp1*-deleted mice are newborn cells. (**A-D**) Control (*Pdgfrα-CreER^TM^*) and *Lrp1*-deleted (*Pdgfrα-CreER^TM^ :: Lrp1^fl/fl^*) mice received cuprizone for 5 weeks, and also received EdU for the 3 finals weeks. Compressed z-stack confocal images show the corpus callosum of control (a, low magnification; b, high magnification) and *Lrp1*-deleted (c, low magnification; d, high magnification) mice labelled to detect the transcription factor OLIG2 (blue) and EdU (red). The vast majority of OLIG2^+^ cells in the corpus callosum of control (146 of 154 cells counted) and *Lrp1*-deleted mice (97 of 106 cells counted) were EdU^+^. Solid magenta arrowheads indicate example OLIG2^+^ EdU^+^ newborn cells. Solid white arrowheads indicate OLIG2^+^ EdU-neg cells. (**E-F**) Compressed z-stack confocal images of the corpus callosum of cuprizone-fed control (*Pdgfrα-CreER^TM^ :: Rosa26-YFP*) and *Lrp1*-deleted (*Pdgfrα-CreER^TM^ :: Rosa26-YFP :: Lrp1^fl/fl^*) mice immunolabelled to detect PDGFRα (red), YFP (green) and LRP1 (blue). YFP^+^ PDGFRα^+^ parenchymal OPCs in *Lrp1*-deleted mice lacked LRP1 (124 of 124 cells counted), however, the YFP-negative PDGFRα^+^ neural stem cell-derived OPCs had intact LRP1 expression (42 of 42 cells counted). Solid yellow arrowheads indicate YFP^+^ LRP1^+^ PDGFRα^+^ parenchymal OPCs in control tissue. Yellow arrows indicate YFP^+^ LRPI-negative PDGFRαΛ parenchymal OPCs in *Lrp1*-deleted tissue. Solid white arrowheads indicate YFP-neg LRP1^+^ PDGFRα^+^ neural stem cell-derived OPCs. White arrows indicate YFP^+^ LRP1-neg PDGFRα-neg newborn OLs in control and *Lrp1*-deleted tissue. Scale bars represent 34μm (A, C and E-H) or 20μm (B, D). CC = corpus callosum.

## References

1. Pepper RE, Pitman KA, Cullen CL, Young KM. How Do Cells of the Oligodendrocyte Lineage Affect Neuronal Circuits to Influence Motor Function, Memory and Mood? Front Cell Neurosci. Frontiers; 2018;12:399.

2. Rivers LE, Young KM, Rizzi M, Jamen F, Psachoulia K, Wade A, et al. PDGFRA/NG2 glia generate myelinating oligodendrocytes and piriform projection neurons in adult mice. Nature Publishing Group. Nature Publishing Group; 2008 Dec;11(12):1392–401.

3. Dimou L, Simon C, Kirchhoff F, Takebayashi H, Gotz M. Progeny of Olig2-Expressing Progenitors in the Gray and White Matter of the Adult Mouse Cerebral Cortex. Journal of Neuroscience. 2008 Oct 8;28(41):10434–42.

4. Young KM, Psachoulia K, Tripathi RB, Dunn S-J, Cossell L, Attwell D, et al. Oligodendrocyte Dynamics in the Healthy Adult CNS: Evidence for Myelin Remodeling. Neuron. Elsevier Inc; 2013 Mar 6;77(5):873–85.

5. Kang SH, Fukaya M, Yang JK, Rothstein JD, Bergles DE. NG2+ CNS Glial Progenitors Remain Committed to the Oligodendrocyte Lineage in Postnatal Life and following Neurodegeneration. Neuron. Elsevier Inc; 2010 Nov 18;68(4):668–81.

6. Hughes EG, Kang SH, Fukaya M, Bergles DE. Oligodendrocyte progenitors balance growth with self-repulsion to achieve homeostasis in the adult brain. Nature Publishing Group. Nature Publishing Group; 2013 Apr 28;16(6):668–76.

7. Hill RA, Li AM, Grutzendler J. Lifelong cortical myelin plasticity and age-related degeneration in the live mammalian brain. Nature Publishing Group. Nature Publishing Group; 2018 May;21(5):683–95.

8. Zhu X, Bergles DE, Nishiyama A. NG2 cells generate both oligodendrocytes and gray matter astrocytes. Development. 2008;135(1):145–57.

9. Hughes EG, Orthmann-Murphy JL, Langseth AJ, Bergles DE. Myelin remodeling through experience-dependent oligodendrogenesis in the adult somatosensory cortex. Nature Publishing Group. Nature Publishing Group; 2018 May;21(5):696–706.

10. Zhang Y, Argaw AT, Gurfein BT, Zameer A, Snyder BJ, Ge C, et al. Notch1 signaling plays a role in regulating precursor differentiation during CNS remyelination. Proc Natl Acad Sci USA. 2009;106(45):19162–7.

11. Givogri MI, Costa RM, Schonmann V, Silva AJ, Campagnoni AT, Bongarzone ER. Central nervous system myelination in mice with deficient expression of notch1 receptor. Journal of Neuroscience Research. 2002;67(3):309–20.

12. Genoud S, Lappe-Siefke C, Goebbels S, Radtke F, Aguet M, Scherer SS, et al. Notch1 control of oligodendrocyte differentiation in the spinal cord. J Cell Biol. 2002;158(4):709–18.

13. Zhou Y-X, Flint NC, Murtie JC, Le TQ, Armstrong RC. Retroviral lineage analysis of fibroblast growth factor receptor signaling in FGF2 inhibition of oligodendrocyte progenitor differentiation. Glia. John Wiley & Sons, Ltd; 2006;54(6):578–90.

14. Murtie JC, Zhou YX, Le TQ, Armstrong RC. In vivo analysis of oligodendrocyte lineage development in postnatal FGF2 null nice. Glia. 2005 Mar;49(4):542–54.

15. Murcia-Belmonte V, Medina-Rodríguez EM, Bribián A, de Castro F, Esteban PF. ERK1/2 signaling is essential for the chemoattraction exerted by human FGF2 and human anosmin-1 on newborn rat and mouse OPCs via FGFR1. Glia. 2014 Mar;62(3):374–86.

16. Grier MD, West KL, Kelm ND, Fu C, Does MD, Parker B, et al. Loss of mTORC2 signaling in oligodendrocyte precursor cells delays myelination. de Castro F, editor. PLoS ONE. 2017;12(11).

17. Zou Y, Jiang W, Wang J, Li Z, Zhang J, Bu J, et al. Oligodendrocyte Precursor Cell-Intrinsic Effect of Rheb1 Controls Differentiation and Mediates mTORC1-Dependent Myelination in Brain. Journal of Neuroscience. Society for Neuroscience; 2014;34(47):15764–78.

18. Jiang M, Liu L, He X, Wang H, Lin W, Wang H, et al. Regulation of PERK-eIF2α signalling by tuberous sclerosis complex-1 controls homoeostasis and survival of myelinating oligodendrocytes. Nature Communications. Nature Publishing Group; 2016 Jul 15;7(1):12185.

19. McKinnon RD. PDGF -Receptor Signal Strength Controls an RTK Rheostat That Integrates Phosphoinositol 3’-Kinase and Phospholipase C Pathways during Oligodendrocyte Maturation. Journal of Neuroscience. 2005 Apr 6;25(14):3499–508.

20. Rajasekharan S. Intracellular signaling mechanisms directing oligodendrocyte precursor cell migration. J Neurosci. Society for Neuroscience; 2008 Dec 10;28(50):13365–7.

21. Chew L-J, Coley W, Cheng Y, Gallo V. Mechanisms of regulation of oligodendrocyte development by p38 mitogen-activated protein kinase. J Neurosci. Society for Neuroscience; 2010 Aug 18;30(33):11011–27.

22. Cahoy JD, Emery B, Kaushal A, Foo LC, Zamanian JL, Christopherson KS, et al. A Transcriptome Database for Astrocytes, Neurons, and Oligodendrocytes: A New Resource for Understanding Brain Development and Function. Journal of Neuroscience. 2008 Jan 2;28(1):264–78.

23. Zhang Y, Chen K, Sloan SA, Bennett ML, Scholze AR, O’Keeffe S, et al. An RNA-Sequencing Transcriptome and Splicing Database of Glia, Neurons, and Vascular Cells of the Cerebral Cortex. Journal of Neuroscience. 2014 Sep 3;34(36):11929–47.

24. Hrvatin S, Hochbaum DR, Nagy MA, Cicconet M, Robertson K, Cheadle L, et al. Single-cell analysis of experience-dependent transcriptomic states in the mouse visual cortex. Nature Publishing Group. Nature Publishing Group; 2018 Jan;21(1):120–9.

25. Auderset L, Cullen CL, Young KM. Low Density Lipoprotein-Receptor Related Protein 1 Is Differentially Expressed by Neuronal and Glial Populations in the Developing and Mature Mouse Central Nervous System. Coulson EJ, editor. PLoS ONE. Public Library of Science; 2016;11(6):e0155878–22.

26. Auderset L, Landowski LM, Foa L, Young KM. Low Density Lipoprotein Receptor Related Proteins as Regulators of Neural Stem and Progenitor Cell Function. Stem Cells International. 2016;2016(4746):2108495–16.

27. Bres EE, Faissner A. Low Density Receptor-Related Protein 1 Interactions With the Extracellular Matrix: More Than Meets the Eye. Front Cell Dev Biol. Frontiers; 2019 Mar 15;7:1916.

28. Parkyn CJ, Vermeulen EGM, Mootoosamy RC, Sunyach C, Jacobsen C, Oxvig C, et al. LRP1 controls biosynthetic and endocytic trafficking of neuronal prion protein. J Cell Sci. 2008 Feb 26;121(6):773–83.

29. Van Gool B, Storck SE, Reekmans SM, Lechat B, Gordts PLSM, Pradier L, et al. LRP1 Has a Predominant Role in Production over Clearance of Aβ in a Mouse Model of Alzheimer’s Disease. Mol Neurobiol. Springer US; 2019 Apr 19;330(6012):1774–12.

30. Liu C-C, Hu J, Zhao N, Wang J, Na W, Cirrito JR, et al. Astrocytic LRP1 Mediates Brain Aβ Clearance and Impacts Amyloid Deposition. J Neurosci. 2017 Mar 8;37(15):3442–16–4031.

31. Cam JA, Zerbinatti CV, Li Y, Bu G. Rapid Endocytosis of the Low Density Lipoprotein Receptor-related Protein Modulates Cell Surface Distribution and Processing of the -Amyloid Precursor Protein. Journal of Biological Chemistry. 2005 Apr 8;280(15):15464–70.

32. Yamamoto K, Owen K, Parker AE, Scilabra SD, Dudhia J, Strickland DK, et al. Low density lipoprotein receptor-related protein 1 (LRP1)-mediated endocytic clearance of a disintegrin and metalloproteinase with thrombospondin motifs-4 (ADAMTS-4): functional differences of non-catalytic domains of ADAMTS-4 and ADAMTS-5 in LRP1 binding. J Biol Chem. 2014 Mar 7;289(10):6462–74.

33. Su EJ, Fredriksson L, Geyer M, Folestad E, Cale J, Andrae J, et al. Activation of PDGF-CC by tissue plasminogen activator impairs blood-brain barrier integrity during ischemic stroke. Nature Medicine. 2008 Jul;14(7):731–7.

34. Rauch JN, Luna G, Guzman E, Audouard M, Challis C, Sibih YE, et al. LRP1 is a master regulator of tau uptake and spread. Nature. 2020 Apr;580(7803):381–5.

35. Kadurin I, Rothwell SW, Lana B, Nieto-Rostro M, Dolphin AC. LRP1 influences trafficking of N-type calcium channels via interaction with the auxiliary α2δ-1 subunit. Sci Rep. 2017 Mar 3;7:43802.

36. Maier W, Bednorz M, Meister S, Roebroek A, Weggen S, Schmitt U, et al. LRP1 is critical for the surface distribution and internalization of the NR2B NMDA receptor subtype. Neuronal LRP1 Regulates Glucose Metabolism and Insulin Signaling in the Brain. Journal of Neuroscience. 2015 Apr 8;35(14):5851–9.

38. Wujak L, Böttcher RT, Pak O, Frey H, Agha El E, Chen Y, et al. Low density lipoprotein receptor-related protein 1 couples β1 integrin activation to degradation. Cell Mol Life Sci. Springer International Publishing; 2018 May;75(9):1671–85.

39. Boyé K, Pujol N, D Alves I, Chen Y-P, Daubon T, Lee Y-Z, et al. The role of CXCR3/LRP1 cross-talk in the invasion of primary brain tumors. Nature Communications. 2017 Nov 17;8(1):1571.

40. Nakajima C, Kulik A, Frotscher M, Herz J, Schäfer M, Bock HH, et al. Low Density Lipoprotein Receptor-related Protein 1 (LRP1) Modulates N-Methyl-d-aspartate (NMDA) Receptor-dependent Intracellular Signaling and NMDA-induced Regulation of Postsynaptic Protein Complexes. Journal of Biological Chemistry. 2013 Jul 26;288(30):21909–23.

41. Polavarapu R, Gongora MC, Yi H, Ranganthan S, Lawrence DA, Strickland D, et al. Tissue-type plasminogen activator-mediated shedding of astrocytic low-density lipoprotein receptor-related protein increases the permeability of the neurovascular unit. Blood. 2007 Apr 15;109(8):3270–8.

42. Brifault C, Gilder AS, Laudati E, Banki M, Gonias SL. Shedding of membrane-associated LDL receptor-related protein-1 from microglia amplifies and sustains neuroinflammation. Journal of Biological Chemistry. American Society for Biochemistry and Molecular Biology; 2017;292(45):18699–712.

43. Brifault C, Kwon H, Campana WM, Gonias SL. LRP1 deficiency in microglia blocks neuro-inflammation in the spinal dorsal horn and neuropathic pain processing. Glia. 2019 Jun;67(6):1210–24.

44. Actis Dato V, Chiabrando GA. The Role of Low-Density Lipoprotein Receptor-Related Protein 1 in Lipid Metabolism, Glucose Homeostasis and Inflammation. Int J Mol Sci. 2018 Jun 15;19(6):1780.

45. Zhuo M, Holtzman DM, Li Y, Osaka H, DeMaro J, Jacquin M, et al. Role of tissue plasminogen activator receptor LRP in hippocampal long-term potentiation. J Neurosci. 2000 Jan 15;20(2):542–9.

46. Herz J, Clouthier DE, Hammer RE. LDL receptor-related protein internalizes and degrades uPA-PAI-1 complexes and is essential for embryo implantation. Cell. 1992 Oct 30;71(3):411–21.

47. Hennen E, Safina D, Haussmann U, Worsdorfer P, Edenhofer F, Poetsch A, et al. A LewisX Glycoprotein Screen Identifies the Low Density Lipoprotein Receptor-related Protein 1 (LRP1) as a Modulator of Oligodendrogenesis in Mice. Journal of Biological Chemistry. 2013 Jun 7;288(23):16538–45.

48. Safina D, Schlitt F, Romeo R, Pflanzner T, Pietrzik CU, Narayanaswami V, et al. Low-density lipoprotein receptor-related protein 1 is a novel modulator of radial glia stem cell proliferation, survival, and differentiation. Glia. 2016 Aug 1;64(8):1363–80.

49. Lin J-P, Mironova YA, Shrager P, Giger RJ. LRP1 regulates peroxisome biogenesis and cholesterol homeostasis in oligodendrocytes and is required for proper CNS myelin development and repair. Elife. eLife Sciences Publications Limited; 2017 Dec 18;6:e30498.

50. Fernandez-Castaneda A, Chappell MS, Rosen DA, Seki SM, Beiter RM, Johanson DM, et al. The active contribution of OPCs to neuroinflammation is mediated by LRP1. Acta Neuropathol. Springer Berlin Heidelberg; 2019 Sep 24;14:1142–18.

51. Velez-Fort M, Maldonado PP, Butt AM, Audinat E, Angulo MC. Postnatal Switch from Synaptic to Extrasynaptic Transmission between Interneurons and NG2 Cells. Journal of Neuroscience. 2010 May 19;30(20):6921–9.

52. Pitman KA, Ricci R, Gasperini R, Beasley S, Pavez M, Charlesworth J, et al. The voltage-gated calcium channel CaV1.2 promotes adult oligodendrocyte progenitor cell survival in the mouse corpus callosum but not motor cortex. Glia. 2020 Oct 12;22:54.

53. Spitzer SO, Sitnikov S, Kamen Y, Evans KA, Kronenberg-Versteeg D, Dietmann S, et al. Oligodendrocyte Progenitor Cells Become Regionally Diverse and Heterogeneous with Age. Neuron. 2019;101(3):459–.

54. Hamilton TG, Klinghoffer RA, Corrin PD, Soriano P. Evolutionary Divergence of Platelet-Derived Growth Factor Alpha Receptor Signaling Mechanisms. Molecular and Cellular Biology. 2003 Jun 1;23(11):4013–25.

55. Srinivas S, Watanabe T, Lin CS, William CM, Tanabe Y, Jessell TM, et al. Cre reporter strains produced by targeted insertion of EYFP and ECFP into the ROSA26 locus. BMC Dev Biol. 2001;1(1):4.

56. Hippenmeyer S, Vrieseling E, Sigrist M, Portmann T, Laengle C, Ladle DR, et al. A developmental switch in the response of DRG neurons to ETS transcription factor signaling. Sanes JR, editor. PLoS Biol. Public Library of Science; 2005 May;3(5):e159.

57. O’Rourke M, Cullen CL, Auderset L, Pitman KA, Achatz D, Gasperini R, et al. Evaluating Tissue-Specific Recombination in a Pdgfrα-CreERT2 Transgenic Mouse Line. PLoS ONE. 2016;11(9):e0162858.

58. Clarke LE, Young KM, Hamilton NB, Li H, Richardson WD, Attwell D. Properties and fate of oligodendrocyte progenitor cells in the corpus callosum, motor cortex, and piriform cortex of the mouse. J Neurosci. Society for Neuroscience; 2012 Jun 13;32(24):8173–85.

59. Emery B, Dugas JC. Purification of oligodendrocyte lineage cells from mouse cortices by immunopanning. Cold Spring Harb Protoc. Cold Spring Harbor Laboratory Press; 2013 Sep 1;2013(9):854–68.

60. Fulton D, Paez PM, Fisher R, Handley V, Colwell CS, Campagnoni AT. Regulation of L-type Ca++ currents and process morphology in white matter oligodendrocyte precursor cells by golli-myelin proteins. Glia. Wiley Subscription Services, Inc., A Wiley Company; 2010 Aug 15;58(11):1292–303.

61. Haberlandt C, Derouiche A, Wyczynski A, Haseleu J, Pohle J, Karram K, et al. Gray matter NG2 cells display multiple Ca2+-signaling pathways and highly motile processes. Degtyar V, editor. PLoS ONE. 2011 Mar 24;6(3):e17575.

62. Psachoulia K, Jamen F, Young KM, Richardson WD. Cell cycle dynamics of NG2 cells in the postnatal and ageing brain. Neuron Glia Biol. Cambridge University Press; 2009 Nov;5(3-4):57–67.

63. Takayama Y, May P, Anderson RGW, Herz J. Low density lipoprotein receptor-related protein 1 (LRP1) controls endocytosis and c-CBL-mediated ubiquitination of the platelet-derived growth factor receptor beta (PDGFR beta). Journal of Biological Chemistry. 2005 May 6;280(18):18504–10.

64. Pi X, Schmitt CE, Xie L, Portbury AL, Wu Y, Lockyer P, et al. LRP1-dependent endocytic mechanism governs the signaling output of the bmp system in endothelial cells and in angiogenesis. Circ Res. Lippincott Williams & Wilkins Hagerstown, MD; 2012 Aug 17;111(5):564–74.

65. Liu Q, Trotter J, Zhang J, Peters MM, Cheng H, Bao J, et al. Neuronal LRP1 knockout in adult mice leads to impaired brain lipid metabolism and progressive, age-dependent synapse loss and neurodegeneration. J Neurosci. Society for Neuroscience; 2010 Dec 15;30(50):17068–78.

66. Pringle N, Collarini EJ, Mosley MJ, Heldin CH, Westermark B, Richardson WD. PDGF A chain homodimers drive proliferation of bipotential (O-2A) glial progenitor cells in the developing rat optic nerve. EMBO J. European Molecular Biology Organization; 1989 Apr;8(4):1049–56.

67. Richardson WD, Pringle N, Mosley MJ, Westermark B, Dubois-Dalcg M. A role for platelet-derived growth factor in normal gliogenesis in the central nervous system. Cell. 1988 Apr;53(2):309–19.

68. Kukley M, Nishiyama A, Dietrich D. The fate of synaptic input to NG2 glial cells: neurons specifically downregulate transmitter release onto differentiating oligodendroglial cells. J Neurosci. Society for Neuroscience; 2010 Jun 16;30(24):8320–31.

69. De Biase LM, Nishiyama A, Bergles DE. Excitability and Synaptic Communication within the Oligodendrocyte Lineage. Journal of Neuroscience. 2010 Mar 10;30(10):3600–11.

70. Trapp BD, Nishiyama A, Cheng D, Macklin W. Differentiation and death of premyelinating oligodendrocytes in developing rodent brain. J Cell Biol. 1997 Apr 21;137(2):459–68.

71. Tripathi RB, Jackiewicz M, McKenzie IA, Kougioumtzidou E, Grist M, Richardson WD. Remarkable Stability of Myelinating Oligodendrocytes in Mice. CellReports. 2017 Oct 10;21(2):316–23.

72. Gan M, Jiang P, McLean P, Kanekiyo T, Bu G. Low-Density Lipoprotein Receptor-Related Protein 1 (LRP1) Regulates the Stability and Function of GluA1 α-Amino-3-Hydroxy-5-Methyl-4-Isoxazole Propionic Acid (AMPA) Receptor in Neurons. Tang Y-P, editor. PLoS ONE. 2014 Dec 12;9(12):e113237–18.

73. Zonouzi M, Renzi M, Farrant M, Cull-Candy SG. Bidirectional plasticity of calcium-permeable AMPA receptors in oligodendrocyte lineage cells. Nature Publishing Group. Nature Publishing Group; 2011 Oct 9;14(11):1430–8.

74. Gallo V, Zhou JM, McBain CJ, Wright P, Knutson PL, Armstrong RC. Oligodendrocyte progenitor cell proliferation and lineage progression are regulated by glutamate receptor-mediated K+ channel block. Journal of Neuroscience. Society for Neuroscience; 1996 Apr 15;16(8):2659–70.

75. Gudz TI. Glutamate Stimulates Oligodendrocyte Progenitor Migration Mediated via an v Integrin/Myelin Proteolipid Protein Complex. Journal of Neuroscience. 2006 Mar 1;26(9):2458–66.

76. Kougioumtzidou E, Shimizu T, Hamilton NB, Tohyama K, Sprengel R, Monyer H, et al. Signalling through AMPA receptors on oligodendrocyte precursors promotes myelination by enhancing oligodendrocyte survival. Elife. eLife Sciences Publications Limited; 2017 Jun 13;6:31.

77. Fannon J, Tarmier W, Fulton D. Neuronal activity and AMPA-type glutamate receptor activation regulates the morphological development of oligodendrocyte precursor cells. Glia. John Wiley & Sons, Ltd; 2015 Jun;63(6):1021–35.

78. Cheli VT, Santiago González DA, Spreuer V, Paez PM. Voltage-gated Ca2+ entry promotes oligodendrocyte progenitor cell maturation and myelination in vitro. Experimental Neurology. 2015 Mar;265:69–83.

79. Williamson AV, Compston DA, Randall AD. Analysis of the ion channel complement of the rat oligodendrocyte progenitor in a commonly studied in vitro preparation. Eur J Neurosci. 1997 Apr;9(4):706–20.

80. Bu G, Morton PA, Schwartz AL. Identification and partial characterization by chemical cross-linking of a binding protein for tissue-type plasminogen activator (t-PA) on rat hepatoma cells. A plasminogen activator inhibitor type 1-independent t-PA receptor. Journal of Biological Chemistry. 1992 Aug 5;267(22):15595–602.

81. Orth K, Madison EL, Gething MJ, Sambrook JF, Herz J. Complexes of tissue-type plasminogen activator and its serpin inhibitor plasminogen-activator inhibitor type 1 are internalized by means of the low density lipoprotein receptor-related protein/alpha 2-macroglobulin receptor. PNAS. 1992 Aug 15;89(16):7422–6.

82. Noble M, Murray K, Stroobant P, Waterfield MD, Riddle P. Platelet-derived growth factor promotes division and motility and inhibits premature differentiation of the oligodendrocyte/type-2 astrocyte progenitor cell. Nature. Nature Publishing Group; 1988 Jun 9;333(6173):560–2.

83. Muratoglu SC, Mikhailenko I, Newton C, Migliorini M, Strickland DK. Low density lipoprotein receptor-related protein 1 (LRP1) forms a signaling complex with platelet-derived growth factor receptor-beta in endosomes and regulates activation of the MAPK pathway. Journal of Biological Chemistry. American Society for Biochemistry and Molecular Biology; 2010 May 7;285(19):14308–17.

84. Spuch C, Ortolano S, Navarro C. LRP-1 and LRP-2 receptors function in the membrane neuron. Trafficking mechanisms and proteolytic processing in Alzheimer’s disease. Front Physiol. Frontiers; 2012 Jul 16;3.

85. Gajera CR, Emich H, Lioubinski O, Christ A, Beckervordersandforth-Bonk R, Yoshikawa K, et al. LRP2 in ependymal cells regulates BMP signaling in the adult neurogenic niche. J Cell Sci. 2010 Jun 1;123(Pt 11):1922–30.

86. Andersen RK, Hammer K, Hager H, Christensen JN, Ludvigsen M, Honoré B, et al. Melanoma tumors frequently acquire LRP2/megalin expression, which modulates melanoma cell proliferation and survival rates. Pigment Cell Melanoma Res. 2015 May;28(3):267–80.

87. Mao H, Lockyer P, Townley-Tilson WHD, Xie L, Pi X. LRP1 Regulates Retinal Angiogenesis by Inhibiting PARP-1 Activity and Endothelial Cell Proliferation. Arteriosclerosis, Thrombosis, and Vascular Biology. Lippincott Williams & Wilkins Hagerstown, MD; 2016 Feb;36(2):350–60.

88. Lin L, Bu G, Mars WM, Reeves WB, Tanaka S, Hu K. tPA activates LDL receptor-related protein 1-mediated mitogenic signaling involving the p90RSK and GSK3beta pathway. The American Journal of Pathology. 2010 Oct;177(4):1687–96.

89. Xing YL, Röth PT, Stratton JAS, Chuang BHA, Danne J, Ellis SL, et al. Adult neural precursor cells from the subventricular zone contribute significantly to oligodendrocyte regeneration and remyelination. J Neurosci. Society for Neuroscience; 2014 Oct 15;34(42):14128–46.

90. Fard MK, van der Meer F, Sánchez P, Cantuti-Castelvetri L, Mandad S, Jäkel S, et al. BCAS1 expression defines a population of early myelinating oligodendrocytes in multiple sclerosis lesions. Sci Transl Med. 2017 Dec 6;9(419):eaam7816.

91. Ferreira S, Pitman KA, Summers BS, Wang S, Young KM, Cullen CL. Oligodendrogenesis increases in hippocampal grey and white matter prior to locomotor or memory impairment in an adult mouse model of tauopathy. Eur J Neurosci. 2020 Mar 17;:ejn.14726.

92. Lillis AP, Van Duyn LB, Murphy-Ullrich JE, Strickland DK. LDL Receptor-Related Protein 1: Unique Tissue-Specific Functions Revealed by Selective Gene Knockout Studies. Physiological Reviews. 2008 Jul 1;88(3):887–918.

93. Boucher P, Li W-P, Matz RL, Takayama Y, Auwerx J, Anderson RGW, et al. LRP1 Functions as an Atheroprotective Integrator of TGFβ and PDGF Signals in the Vascular Wall: Implications for Marfan Syndrome. PLoS ONE. 2007 May 16;2(5):e448.

94. Mantuano E, Jo M, Gonias SL, Campana WM. Low density lipoprotein receptor-related protein (LRP1) regulates Rac1 and RhoA reciprocally to control Schwann cell adhesion and migration. J Biol Chem. 2010 May 7;285(19):14259–66.

95. Mantuano E, Lam MS, Shibayama M, Campana WM, Gonias SL. The NMDA receptor functions independently and as an LRP1 co-receptor to promote Schwann cell survival and migration. J Cell Sci. The Company of Biologists Ltd; 2015 Sep 15;128(18):3478–88.

96. Barcelona PF, Jaldin-Fincati JR, Sanchez MC, Chiabrando GA. Activated 2-macroglobulin induces Muller glial cell migration by regulating MT1-MMP activity through LRP1. The FASEB Journal. 2013 Aug 1;27(8):3181–97.

97. Sayre N, Kokovay E. Loss of lipoprotein receptor LRP1 ablates neural stem cell migration to ischemic lesions. The FASEB Journal. The Federation of American Societies for Experimental Biology; 2019 Apr 1.

98. Ferrer DG, Actis Dato V, Jaldín Fincati JR, Lorenc VE, Sánchez MC, Chiabrando GA. Activated α2 -Macroglobulin Induces Mesenchymal Cellular Migration of Raw264.7 Cells through Low-Density Lipoprotein Receptor-Related Protein 1. J Cell Biochem. 2016 Dec 24.

99. Zucker MM, Wujak L, Gungl A, Didiasova M, Kosanovic D, Petrovic A, et al. LRP1 promotes synthetic phenotype of pulmonary artery smooth muscle cells in pulmonary hypertension. Biochim Biophys Acta Mol Basis Dis. 2019 Jun 1;1865(6):1604–16.

100. Yang T-H, St John LS, Garber HR, Kerros C, Ruisaard KE, Clise-Dwyer K, et al. Membrane-Associated Proteinase 3 on Granulocytes and Acute Myeloid Leukemia Inhibits T Cell Proliferation. J Immunol. American Association of Immunologists; 2018 Sep 1;201(5):1389–99.

101. Basford JE, Moore ZWQ, Zhou L, Herz J, Hui DY. Smooth muscle LDL receptor-related protein-1 inactivation reduces vascular reactivity and promotes injury-induced neointima formation. Arteriosclerosis, Thrombosis, and Vascular Biology. Lippincott Williams & Wilkins; 2009 Nov;29(11):1772–8.

102. Boucher P, Gotthardt M, Li WP, Anderson R, Herz J. LRP: Role in vascular wall integrity and protection from atherosclerosis. Science. American Association for the Advancement of Science; 2003;300(5617):329–32.

103. Kang SH, Fukaya M, Yang JK, Rothstein JD, Bergles DE. NG2+ CNS Glial Progenitors Remain Committed to the Oligodendrocyte Lineage in Postnatal Life and following Neurodegeneration. Neuron. Elsevier Inc; 2010 Nov 18;68(4):668–81.

104. Zhu X, Hill RA, Dietrich D, Komitova M, Suzuki R, Nishiyama A. Age-dependent fate and lineage restriction of single NG2 cells. Development. 2011 Feb;138(4):745–53.

105. Marshall CAG, Novitch BG, Goldman JE. Olig2 directs astrocyte and oligodendrocyte formation in postnatal subventricular zone cells. Journal of Neuroscience. 2005 Aug;25(32):7289–98.

106. Cai J, Chen Y, Cai W-H, Hurlock EC, Wu H, Kernie SG, et al. A crucial role for Olig2 in white matter astrocyte development. Development. The Company of Biologists Ltd; 2007 May;134(10):1887–99.

107. Xiao L, Ohayon D, McKenzie IA, Sinclair-Wilson A, Wright JL, Fudge AD, et al. Rapid production of new oligodendrocytes is required in the earliest stages of motor-skill learning. Nature Publishing Group. Nature Publishing Group; 2016 Sep;19(9):1210–7.

108. Baxi EG, DeBruin J, Jin J, Strasburger HJ, Smith MD, Orthmann-Murphy JL, et al. Lineage tracing reveals dynamic changes in oligodendrocyte precursor cells following cuprizone-induced demyelination. Glia. John Wiley & Sons, Ltd; 2017 Dec;65(12):2087–98.

109. Miron VE, Kuhlmann T, Antel JP. Cells of the oligodendroglial lineage, myelination, and remyelination. BBA -Molecular Basis of Disease. Elsevier B.V; 2011 Feb 1;1812(2):184–93.

110. Wang C, Zhang C-J, Martin BN, Bulek K, Kang Z, Zhao J, et al. IL-17 induced NOTCH1 activation in oligodendrocyte progenitor cells enhances proliferation and inflammatory gene expression. Nature Communications. Nature Publishing Group; 2017 May 31;8(1):15508–16.

111. Kirby L, Jin J, Cardona JG, Smith MD, Martin KA, Wang J, et al. Oligodendrocyte precursor cells present antigen and are cytotoxic targets in inflammatory demyelination. Nature Communications. Nature Publishing Group; 2019 Aug 29;10(1):3887–20.

112. Falcao AM, van Bruggen D, Marques S, Meijer M, Jäkel S, Agirre E, et al. Disease-specific oligodendrocyte lineage cells arise in multiple sclerosis. Nature Medicine. Nature Publishing Group; 2018 Dec;24(12):1837–44.

113. Fernandez-Castaneda A, Arandjelovic S, Stiles TL, Schlobach RK, Mowen KA, Gonias SL, et al. Identification of the Low Density Lipoprotein (LDL) Receptor-related Protein-1 Interactome in Central Nervous System Myelin Suggests a Role in the Clearance of Necrotic Cell Debris. Journal of Biological Chemistry. 2013 Feb 15;288(7):4538–48.

114. Stiles TL, Dickendesher TL, Gaultier A, Fernandez-Castaneda A, Mantuano E, Giger RJ, et al. LDL receptor-related protein-1 is a sialic-acid-independent receptor for myelin-associated glycoprotein that functions in neurite outgrowth inhibition by MAG and CNS myelin. J Cell Sci. 2013 Jan 1;126(Pt 1):209–20.

115. Gaultier A, Wu X, Le Moan N, Takimoto S, Mukandala G, Akassoglou K, et al. Low-density lipoprotein receptor-related protein 1 is an essential receptor for myelin phagocytosis. J Cell Sci. 2009 Apr 15;122(Pt 8):1155–62.

